# Evaluating environmental and ecological landscape characteristics relevant to urban resilience across gradients of land-sharing-sparing and urbanity

**DOI:** 10.1101/605105

**Authors:** Matthew Dennis, Katherine L. Scaletta, Philip James

## Abstract

Within urban landscape planning, debate continues around the relative merits of land-sparing (compaction) and land-sharing (sprawl) scenarios. Using part of Greater Manchester (UK) as a case-study, we present a landscape approach to mapping green infrastructure and variation in social-ecological-environmental conditions as a function of land sparing and sharing. We do so for the landscape as a whole as well as for areas of high and low urbanity. Results imply potential trade-offs between land-sparing-sharing scenarios relevant to characteristics critical to urban resilience such as landscape connectivity and diversity, air quality, surface temperature, and access to green space. These trade-offs may be particularly complex due to the parallel influence of patch attributes such as land-cover and size and imply that both ecological restoration and spatial planning have a role to play in reconciling tensions between land-sparing and sharing strategies.

## Introduction

The concept of green infrastructure has emerged as a promising framework to understand, manage and enhance the multiple benefits delivered from nature, particularly in highly fragmented landscapes (Benedict and McMahon, 2012). A green infrastructure approach involves optimizing multi-functionality in terms of social, ecological and economic benefits (Mell, 2013) and seeking resilience through landscape diversity, connectivity and micro-climate regulation (Lovell and Taylor, 2013). With the unabated growth of urban areas in terms of population and the associated sprawl of developed areas into the rural hinterland, debates surrounding the optimum spatial configuration on which to base urban planning persist. At the centre of this debate is a tension between the relative social-ecological effects of urban densification (or the so-called compact cities approach) versus urban sprawl. This tension is largely characterized by high versus low population densities and associated housing stock (Couch and Karecha, 2006). In scenarios which involve increased urban densification, questions arise as to how urban spatial planning can ensure the provision of adequate green space cover in order to maintain vital ecosystem services to urban residents.

In recent years, a land-sharing versus land-sparing model, borrowed from landscape ecological studies on the effects of agricultural land-use on biodiversity (Phalan, 2011), has been adopted as a means to explore the influence of urbanization on ecological integrity. The model is particularly useful in the context of urbanization given the parallels that exist between the latter and agriculture-driven land-use change on which the concept was originally founded, namely high levels of local species extinction and ecosystem service degradation (Lin and Fuller, 2013). In an urban context, a land-sparing approach is promoted in cases where non-green land-use is compacted in order to allow for larger patches of green space. This template theoretically favours large public green spaces in favour of smaller private green spaces in the form of domestic gardens (Geschke et al., 2017). Conversely, land-sharing implies the promotion of lower-density development which leads to smaller, more fragmented patches of public green space and greater cover by private domestic gardens. However, this dichotomy of public and private green land-use is still poorly understood from ecological, social and environmental points of view. Moreover, there is, as yet, insufficient evidence that public or private green land-use *per se* promotes either sparing or sharing outcomes. This is in large part due to a low number of empirical studies and poorly conceived representations of urban green infrastructure.

### Conceptualizing green infrastructure for urban land sparing-sharing studies

A key shortcoming of both the conceptualization and spatial representation of green infrastructure in research on urban areas is a consideration of green space from either an anthropocentric point of view (i.e. as land-use or function) or from a physical-ecological point of view (i.e. land-cover). In order to understand the relative benefits of land-sparing versus sharing in urban areas, composite datasets are required that can model spatial variation in public and private land-use in tandem with their respective land-covers. With improved datasets, based on more social-ecological conceptualisations of green infrastructure, ecological and socio-environmental characteristics critical to resilience in urban systems could be effectively modelled. Moreover, the assumptions around the role of public versus private urban green space in promoting sparing and sharing scenarios respectively can also be clarified, which should inform persisting debates within urban planning.

However, despite the need for holistic, integrated conceptualisations of urban landscapes, research on urban land sparing and sharing has largely sought to reduce the complex characteristics of urban areas. For example, studies have typically modelled hypothetical landscapes based on observed patterns of species distribution (Caryl et al., 2016) as a response to broad land-use types such as building density (Soga et al., 2014). In addition, meta-analyses drawing on a range of geographically diverse studies (Stott et al., 2015) have been carried out in order to identify common trends. These reductionist approaches however, have not considered wider social-ecological factors such as landscape connectivity, heterogeneity, overall green cover quantity and quality or other socio-environmental factors such as access to nature, urban cooling or air quality. We argue that a more holistic approach to evaluating urban landscapes is necessary in order to inform planning frameworks that align with UN Sustainable Development Goals. The creation of landscapes that promote human well-being, urban resilience to climate change, and which address inequalities in addition to biodiversity loss, requires a green infrastructure approach which considers a range of social-ecological outcomes (Lovell and Taylor, 2013; Reyers et al., 2013; Schewenius et al., 2014; Ramaswami et al., 2016). Doing so is only achievable through the mapping of whole study areas in sufficient spatial and thematic detail. To our knowledge, no studies on land-sparing-sharing scenarios exist that extensively and accurately characterise urban green infrastructure of whole landscapes. The latter is essential in order to model social, ecological and environmental factors vital to sustainable urban planning. For example, landscape connectivity and heterogeneity are positively linked to both the provision and, in particular, the resilience of ecosystem services (Ahern, 2011; Mitchell et al., 2013) whereas attributes such as core area and primary productivity are likewise important indicators of ecosystem service providing landscapes (Kong et al., 2014; Xu et al., 2016).

Urban landscapes are particularly heterogeneous, however, in terms of land-use and highly fragmented in terms of land-cover and, therefore, present significant challenges for the accurate classification and quantification of green infrastructure components. Recent advances in geographic information and remote sensing applications to the mapping of urban areas, employing high resolution open-source data, have provided an opportunity to improve the situation with regards to the generation of fit-for-purpose urban spatial datasets. Recent work by Dennis et al. (2018) and Haase et al. (2019), for example, have demonstrated how a range of geo-computational techniques can be applied to high resolution remotely sensed data integrating information on land-use and land-cover in order to achieve high levels of integration necessary for studying complex social-ecological landscapes. Such advances present an opportunity to explore associations between spatial configurations of green infrastructure and urban social-ecological outcomes.

### Conceptualizing land-sparing-sharing outcomes within a green infrastructure framework

The consideration of wider characteristics such as overall green cover and quality in urban localities is particularly important if urban studies are to be based on the same robust logic as agriculture-based studies on land-sparing-sharing. The latter are assessed primarily at the level of yield-to-species density performance in order to compare the relative success of sparing-to-sharing scenarios (Phalan, 2018). In urban areas however, the management goal is less clear or, at least, characterised with less consistency. Although housing density provides a useful proxy for level of development in urban environments, this comprises only one type of built infrastructure common in urbanizing landscapes. Sophisticated measures of “yield” from urbanisation, comparable to the use of the term in agricultural land-sparing-sharing models, are not forthcoming. A useful approach is to consider total surface sealing as a measure of overall development and, therefore, as a proxy for services delivered by “grey infrastructure”. The question then, from a land-sparing-sharing perspective, is whether consolidating such grey infrastructure into compact forms for the sake of sparing large undeveloped spaces is preferable to allowing developed areas to spread out in low-density patterns. The latter implies smaller, albeit potentially more numerous patches of green space and represents a lower level of urban land-use intensity that, in both agricultural and urbanisation contexts, inevitably requires a larger spatial extent (Stott et al., 2015). However, in the urban context, where measuring productivity is a more complex issue, in order to assess the relative performance of land that remains undeveloped, it is logical to standardise comparisons of land-sharing and land-sparing scenarios by the degree of development and scale. The former requires that, for the same degree of urban development (i.e. surface sealing) a direct comparison across a range of desirable landscape attributes can be made between different spatial configurations. This is important for three reasons. Firstly, without this standardised approach, it is not possible to assess whether relative gains (e.g. land-cover diversity and connectivity) are due to spatial factors or simply a greater amount of green land-cover. Secondly, by taking a standardised approach, meaningful comparisons across scales of investigation are thereby permitted. By developing assessments which model outcomes across scales and are standardised by area, a more informed view can be taken on spatial planning approaches which balance land-use productivity with landscape resilience. Thirdly, decision-makers are required to develop urban spatial frameworks within defined spatial extents according to administrative boundaries. Therefore, research which can identify optimum landscape configurations for a given degree of development at a range of scales are desperately needed in order to allow planners to design urban areas which can provide much needed ecosystem services to local residents. Such knowledge may assist decision-makers to identify bottom lines for the amount of green infrastructure cover necessary at a range of scales that, when consisting of suitable type and distribution, ensures both productivity and resilience.

Land itself can be thought of as the primary asset to be managed in urban areas with local planning authorities working to tight spatial and regulatory constraints, and within administrative boundaries. In light of increasing land-use pressures associated with highly modified urban landscapes, integrated analyses on the relative benefits associated with different landscape patterns are necessary for planners and developers to navigate such complexity. There is a need, therefore, to develop assessments of the relative social, ecological and environmental merits of different urban landscape configurations at meaningful scales (i.e. that are both appropriate to urban governance and transferable between scientific disciplines). Such cross-scale comparisons can only be carried out if whole-landscape studies are facilitated by accurate, integrated characterisations of land-use-land-cover combinations in existing urban landscapes.

### The urban-to-peri-urban context

The spatial and temporal heterogeneity of landscapes subject to urbanisation stand in contrast to the relatively homogenizing effect of land-use change by agriculture and reinforce the need for high resolution, integrated data on urban spatial configurations. Gradients of urbanisation in particular create complex social-ecological conditions. Rural to urban gradients have been shown to exhibit considerable variation in ecosystem service provision (Radford and James, 2010; Haase, 2019), well-being effects of green space (Dennis and James, 2017) and biodiversity outcomes (Turrini and Knop, 2015). Moreover, urbanised landscapes covering city-regions may encompass a range of human-dominated land-uses including highly compacted urban centres to low-density suburbs as well as agricultural landscapes in the peri-urban fringe. Due to such contrasting land-use-land-cover configurations, calls have rightly been made to employ whole-landscape approaches to modelling sparing-sharing outcomes in urban systems (Lin and Fuller, 2013). In addition to whole-landscape assessments we also argue that analyses at sub-landscape scales e.g. within urban and peri-urban zones are necessary given that the subject of a land-sparing-sharing model (i.e. the land being “spared”) will differ depending on the context. For example, taking a sparing approach in high-urban areas will typically imply the promotion of urban intensification towards consolidating larger patches of urban green space whereas, in peri-urban areas, the “spared” land will likely take the form of agricultural or forestry land. This raises another important point related to a land-sharing-sparing dichotomy within the context of urbanisation. Much of the debate and associated research related to land-sparing and sharing in agricultural landscapes is predicated on the relative success of modelled yield-species density curves within biodiversity supporting habitats. However, many peri-urban landscapes typically comprise already degraded ecosystems in various stages of agricultural land-use. Indeed, for some functional groups, urban areas, and residential gardens in particular, can contain higher diversity and abundance than the agricultural hinterland (Cussans et al., 2010). Therefore, it is entirely possible that assumptions applied to land-sparing conservation efforts in areas containing intact biodiversity-supporting vegetation, may not be applicable to landscapes made up of complex juxtapositions of highly-modified land-uses. Given the variance in green infrastructure function, heterogeneity and quality between urban and peri-urban areas, information on vegetation type and health is a critical factor (along with spatial characteristics such as connectivity and patch size) when judging the productivity and resilience of landscapes characterised by (semi-)natural and highly modified habitats.

In order to address these research imperatives, a novel spatial dataset was created, following a method developed by Dennis et al. (2018), which allowed the precise measurement of land-use-land-cover configurations across a spatially contiguous urban area comprising the two cities of Manchester and Salford, and the Metropolitan Borough of Trafford, all parts of Greater Manchester, in north-west England, UK. Our overall aim was to evaluate associations between sharing-sparing scenarios on a range of social-ecological-environmental factors relevant to urban landscape productivity and resilience. In order to do this robustly we focussed on potential mediating factors and, as such, our objectives were three-fold: 1: to assess the relative contribution of land-use-land-cover combinations to sparing-sharing configurations; 2: to identify scale-effects in the performance of sparing-sharing scenarios, and 3: to evaluate the relevance of urban and peri-urban contexts in assessing the relative merits of different landscape configurations.

## Methods

### Spatial data on land-use and land-cover

A composite spatial dataset covering the contiguous urban areas of three districts in Greater Manchester (the cities of Manchester, Salford and the metropolitan borough of Trafford) was achieved through a combination of remote sensing and GIS techniques based on a method published by Dennis et al. (2018). Briefly, the method achieves the characterisation of discrete landscape features through an integration of land-use and land-cover data. Land-use (from OS Mastermap Topography and Greenspace layers, 2017 and UK Land Cover Map: Rowland et al., 2015) was computed for public (including all public parks and amenity green spaces), domestic green space (private gardens including rented allotment gardens), urban fabric, informal urban greenery (street-scapes and informal and/or spontaneous vegetation within the urban fabric), institutional land and peri-urban land-use within the study area. In addition, spatially co-incident data on land-cover were classified through Planet Scope 3 m imagery (Planet Team, 2017) and supplemented by Ordnance Survey Rivers, Woodland and Buildings layers (OS Open Rivers 2018; OS Open Map Local, 2018) and City of Trees canopy data (Cityoftrees.org.uk, 2011), resulting in five classes (built, ground vegetation, field layer vegetation, tree canopy and water). Accuracy assessment of the land-cover layer was achieved through 200 randomly generated sampling points (40 for each land cover type) for which classified values were cross-tabulated with ground truth evaluations using aerial photography (Edina, 2017). Overall accuracy and Cohen’s Kappa co-efficient were subsequently calculated. The work flow for the land-cover classification is summarised in Figure 1.

**Figure 1.**
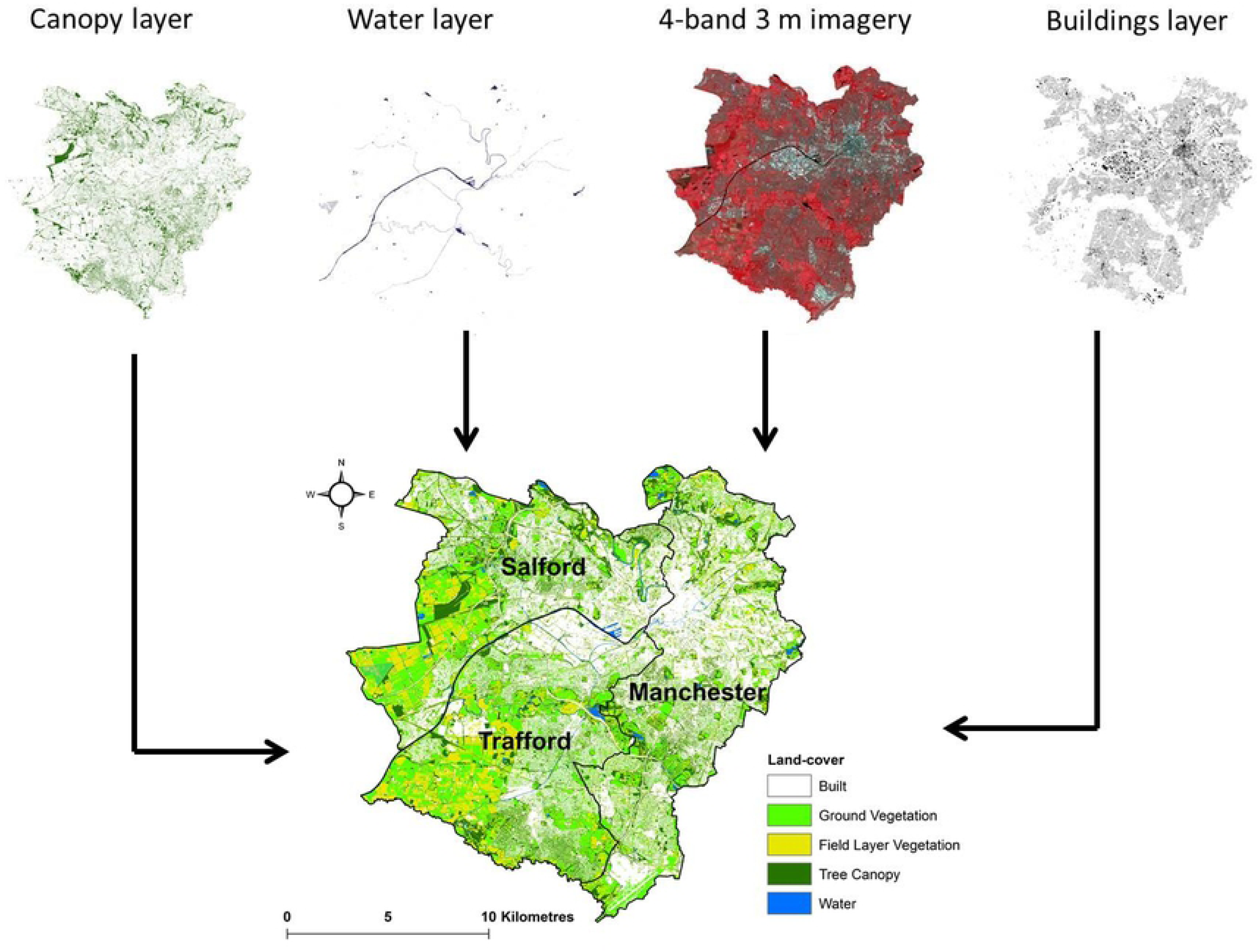
Work-flow for the land-cover classification used in this study combining 3 m satellite imagery (Planet Scope, 2018), tree canopy data (City of Trees 2011 and Ordnance Survey Open Map Local, 2018) and buildings data (OS Open Map Local, 2018).

### Landscape and environmental metrics

A range of social-ecological metrics were quantified within 0.5, 1 and 2 km^2^ zones created through a hexagonal tessellation of the study area. The land-cover layer was used to compute a range of landscape characteristics including effective mesh size (Meff), total core area (TCA), largest patch index (LPI) and Shannon’s land-cover diversity (SHDI), calculated using the QGIS plug-in Lecos (Jung, 2015). Values for Meff and TCA are returned in the spatial units of the source data and, in order to allow comparability across scales, these were standardized as a percentage of the spatial units used in our analysis. In addition, socio-environmental variables land surface temperature (LST, derived from Landsat 8 TIRS imagery for July 2018 at 30 m resolution: NASA, 2018), background nitrogen dioxide concentration (interpolated using the ordinary kriging method from Defra background nitrogen dioxide data points, 2018) and population within 300 m of a recreational green space (using PopGrid 10 m data: Murdock et al., 2017) As a measure of vegetation quality, the normalized difference vegetation index (NDVI) was calculated for pixels in the dataset classified as vegetation (i.e. ground layer, field layer and tree canopy). This was achieved by creating a mask based on all green land-cover pixels and setting this as the environment for the NDVI calculation within ArcMap (version 10.4), again at units of 0.5, 1 and 2 km^2^. We refer to this metric as vNDVI in subsequent sections. Subsequently, the degree to which the tessellated regions exhibited land-cover indicative of land-sparing or land-sharing was judged according to their largest patch index (LPI), following similar approaches taken elsewhere (e.g. Soga et al. 2015). This metric represents the proportion of green space in a given locality that is comprised of a single contiguous patch. High values therefore represent increasing sparing of large patches relative to overall cover by green-space. Tessellated regions were divided into three quantile groups representing low (land-sharing), medium (neither land-sparing nor land sharing) and high (land-sparing) scores for LPI. Figure 2 gives examples of areas exhibiting low, medium and high LPI (land-sharing, neither sharing nor sparing, and land-sparing respectively).

**Figure 2.**
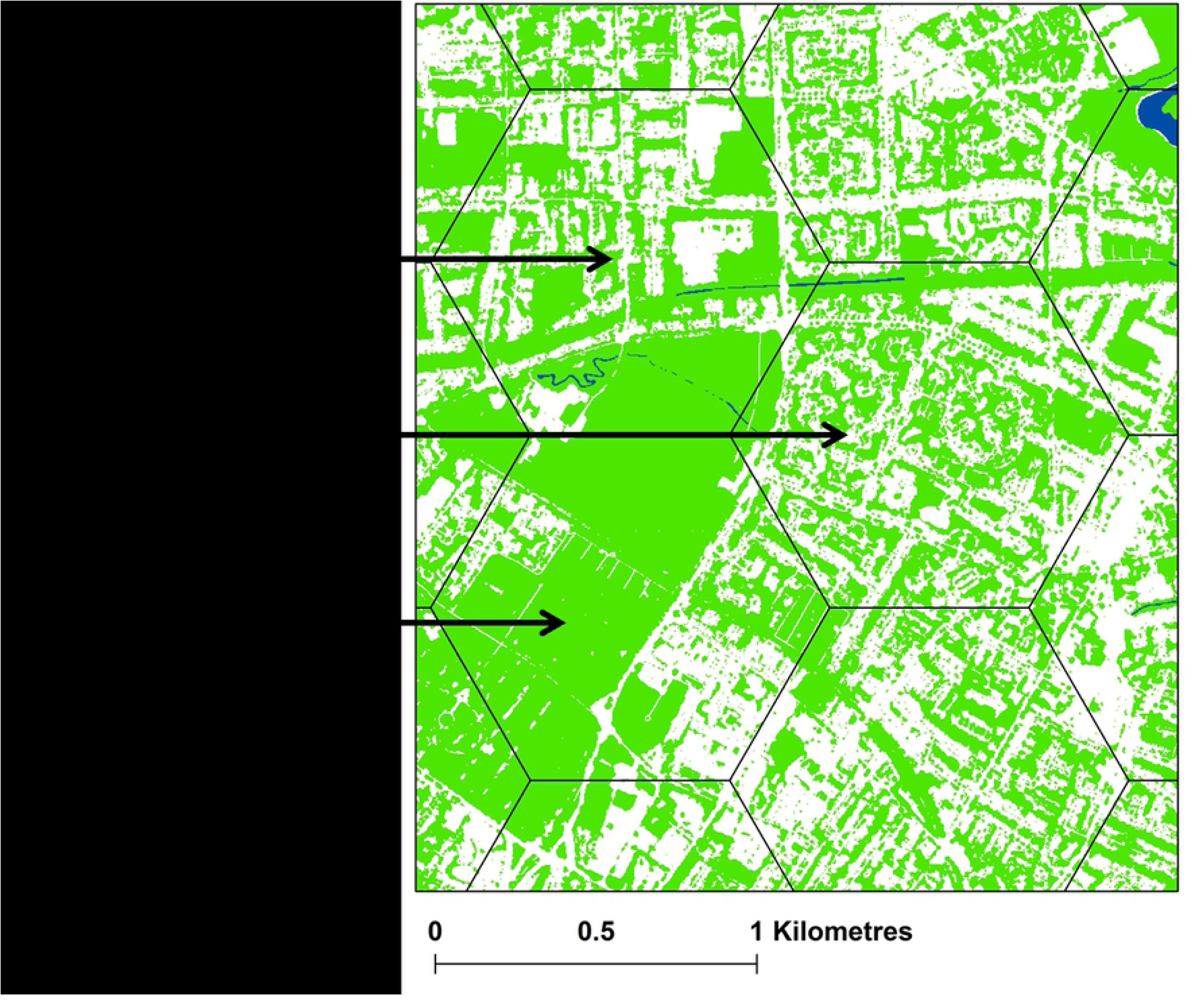
Example of areas classified as land-sharing, land-sparing and neither sharing nor sparing.

The influence of land-sharing/sparing on critical ecological and socio-environmental attributes was assessed through a series of general linear models using the three LPI quantile groups as fixed factors. Meff, SHDI, TCA, vNDVI, LST, nitrogen dioxide and percentage of the local population within 300 m of a recreational green space were all entered as dependent variables whilst controlling for total green land-cover. Controlling for overall green cover, in addition to fulfilling the standardised approach argued for in the introduction to this paper, was equally important from a methodological point of view. LPI and total green land-cover were significantly correlated (at units of 1 km^3^, for example, Pearson’s r = 0.82; p < 0.01). Therefore, entering green land-cover as a co-variate ensured that the LPI metric was not acting as a surrogate for the former in our assessments. Analyses were repeated at low and high urbanity levels (separated by the median values of developed land – i.e. non-green land-use - within each of the 0.5, 1 and 2 km^2^ units of analysis).

Given that socio-economic status is known to influence green cover in urban land-uses (Baker et al., 2018; Dennis et al., 2018) and that the latter may influence the performance of sparing-sharing patterns of green infrastructure, information on vegetation cover within green land-uses was calculated for low and high-urban areas. Income deprivation scores from the English Indices of Multiple Deprivation (DCLG, 2015) were downloaded for Lower Super Output Areas (LSOAs; English census reporting units – mean population is 1500) and mean values were assigned to the smallest unit of analysis for this study (0.5 km^2^ zones; N = 554) in order to best reflect the spatial variance in the original LSOAs dataset (N = 570; mean area = 0.56 km^2^). Finally, associations between land-use-land-cover metrics were explored through multiple linear regression analysis. LPI, TCA, Meff, SHDI, mean LST, mean nitrogen dioxide and mean vNDVI values were entered as dependent variables. The list of land-use-land-cover metrics computed and entered into regression models as independent variables is given in Table 1.

**Table 1.**
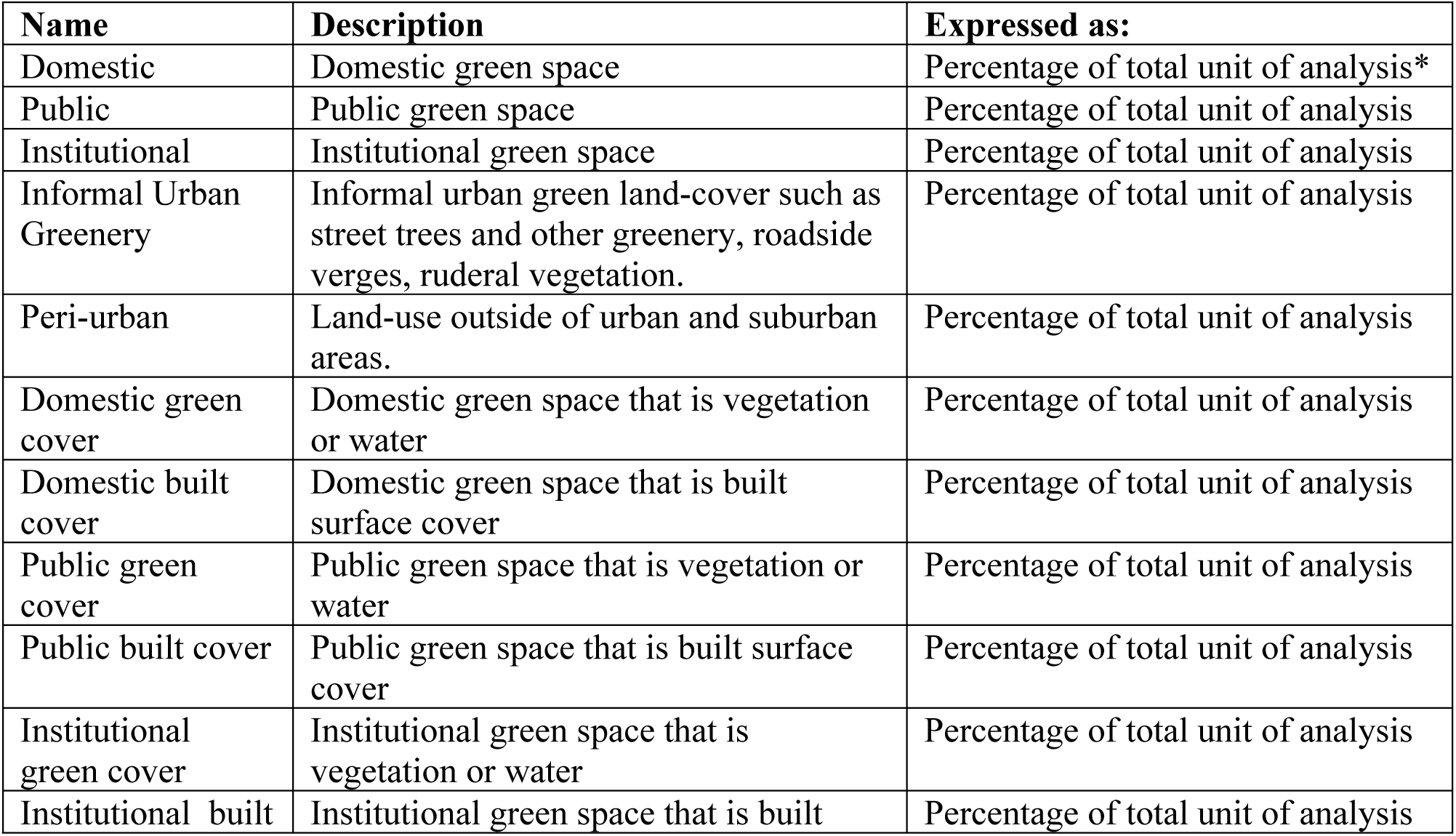
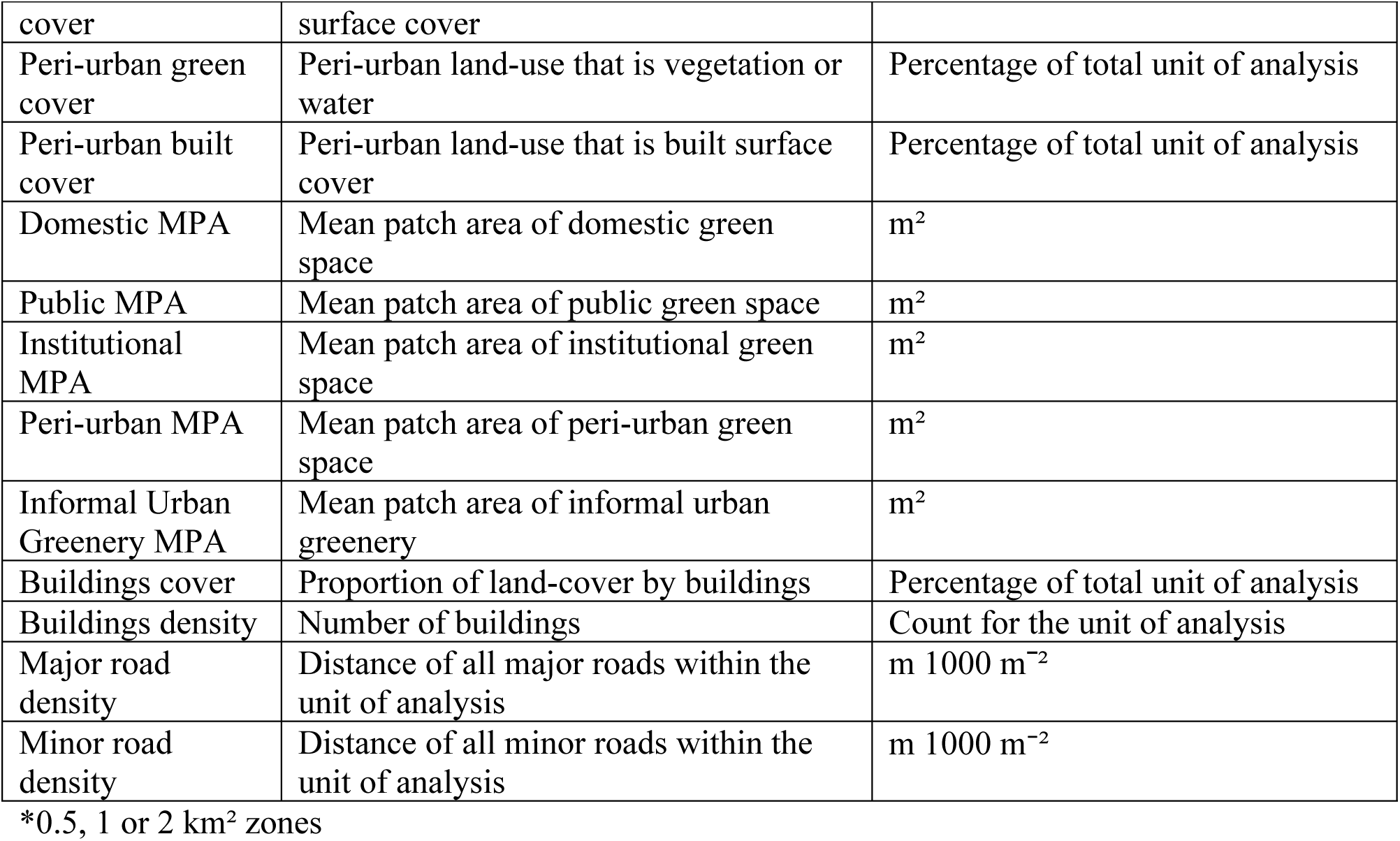
Descriptions of landscape metrics computed for use in linear regression analyses within this study.

In addition to the above, for models in which vegetation type was deemed to be of particular relevance (i.e. where mean LST, nitrogen dioxide and vNDVI were the dependent variables), combinations of all land-use and land-cover classes (proportion of the unit of analysis that is e.g. tree canopy in public parks or ground layer vegetation in the urban fabric) were entered as independent variables. For analyses with mean nitrogen dioxide as the dependent variable, density (m 1000 m^-2^) of major and minor roads (downloaded from OS Open Roads, 2018), were also considered important predictors, as primary emission sources. Regression models were carried out at the 1 km^2^ level as this provided a more robust number of cases than doing so at the 2 km^2^ level whereas an unsatisfactorily high number of missing values for the variables given in Table 1 were produced when calculated at the 0.5 km^2^ level. All statistical tests were carried out in SPSS.23.

## Results

Land-cover and land-use attributes for the study area (form and function) are presented in Figures 3 and 4 respectively. The land-use classification achieved a high level of overall accuracy (92%; Cohen’s Kappa = 0.89, p < 0.001). Figure 5 gives the relative cover by major land-uses (those comprising > 1% of the study area) and associated land-cover across low, medium and high income levels (for whole-landscape and for low versus high-urban areas) at the 0.5 km^2^ level.

**Figure 3.**
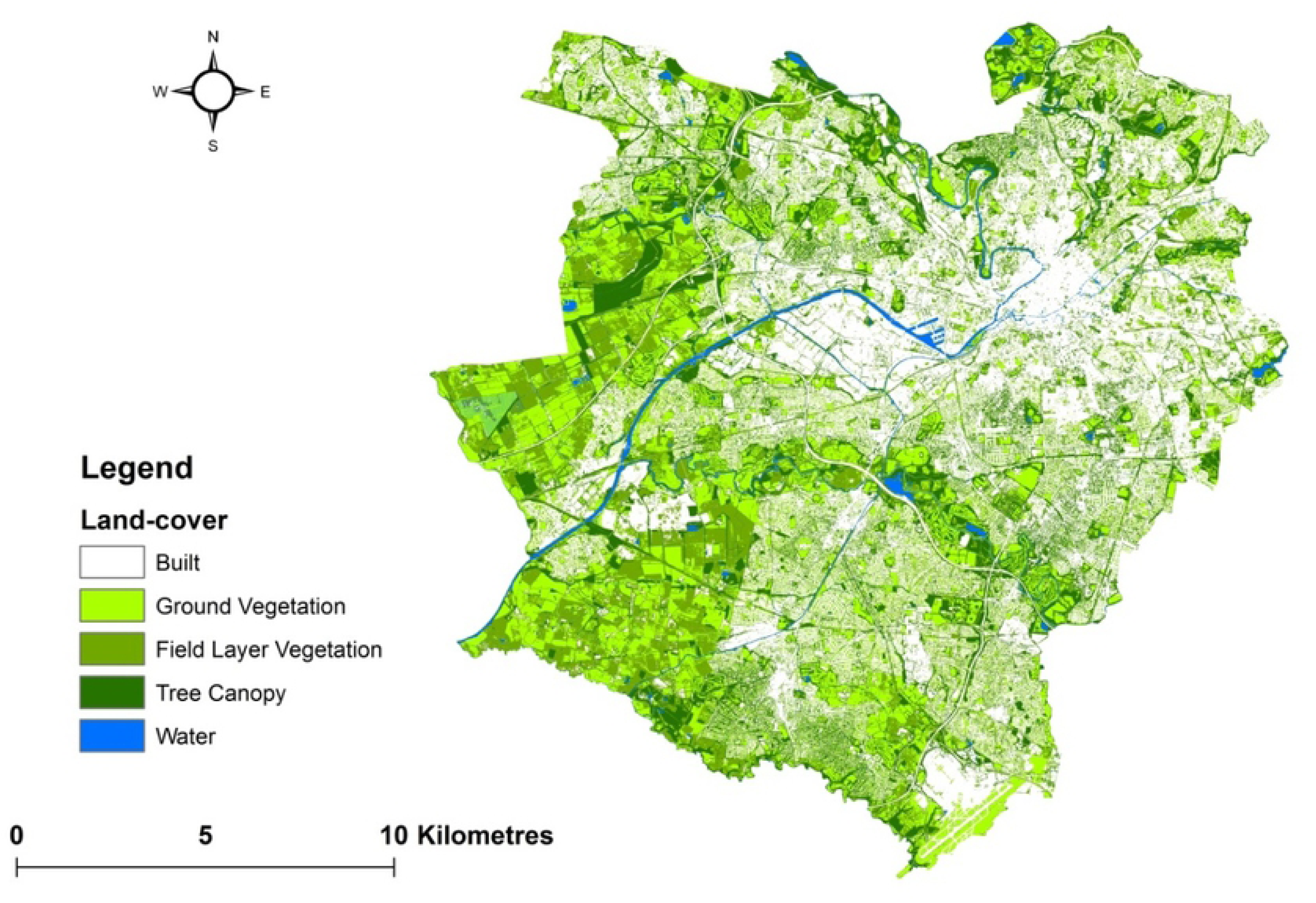
Study area characterised by land-cover (contains Planet Scope 2017, City of Trees 2011 and Ordnance Survey, 2018 data)

**Figure 4.**
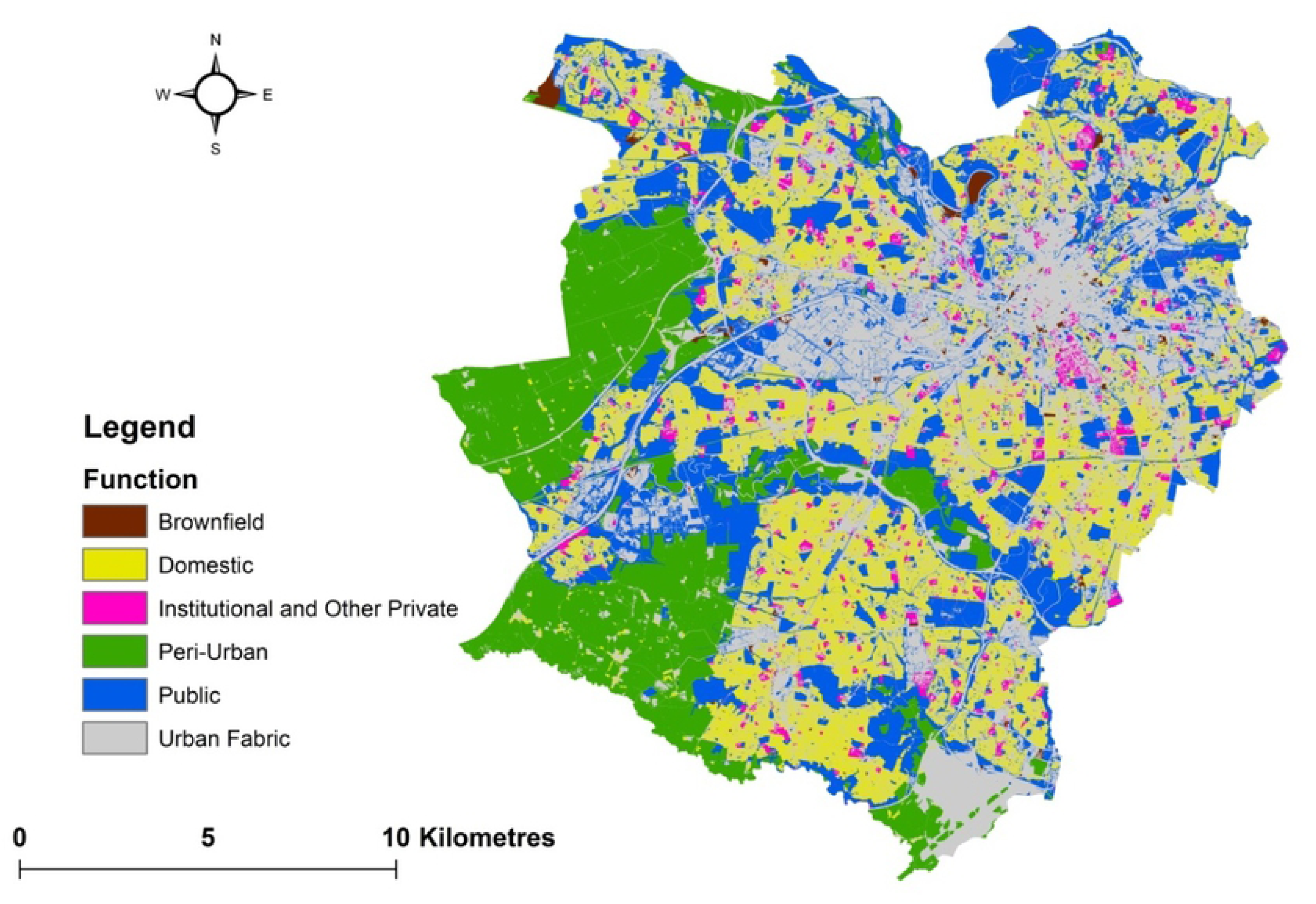
Land-uses within the study area (contains Ordnance Survey 2018 data)

**Figure 5.**
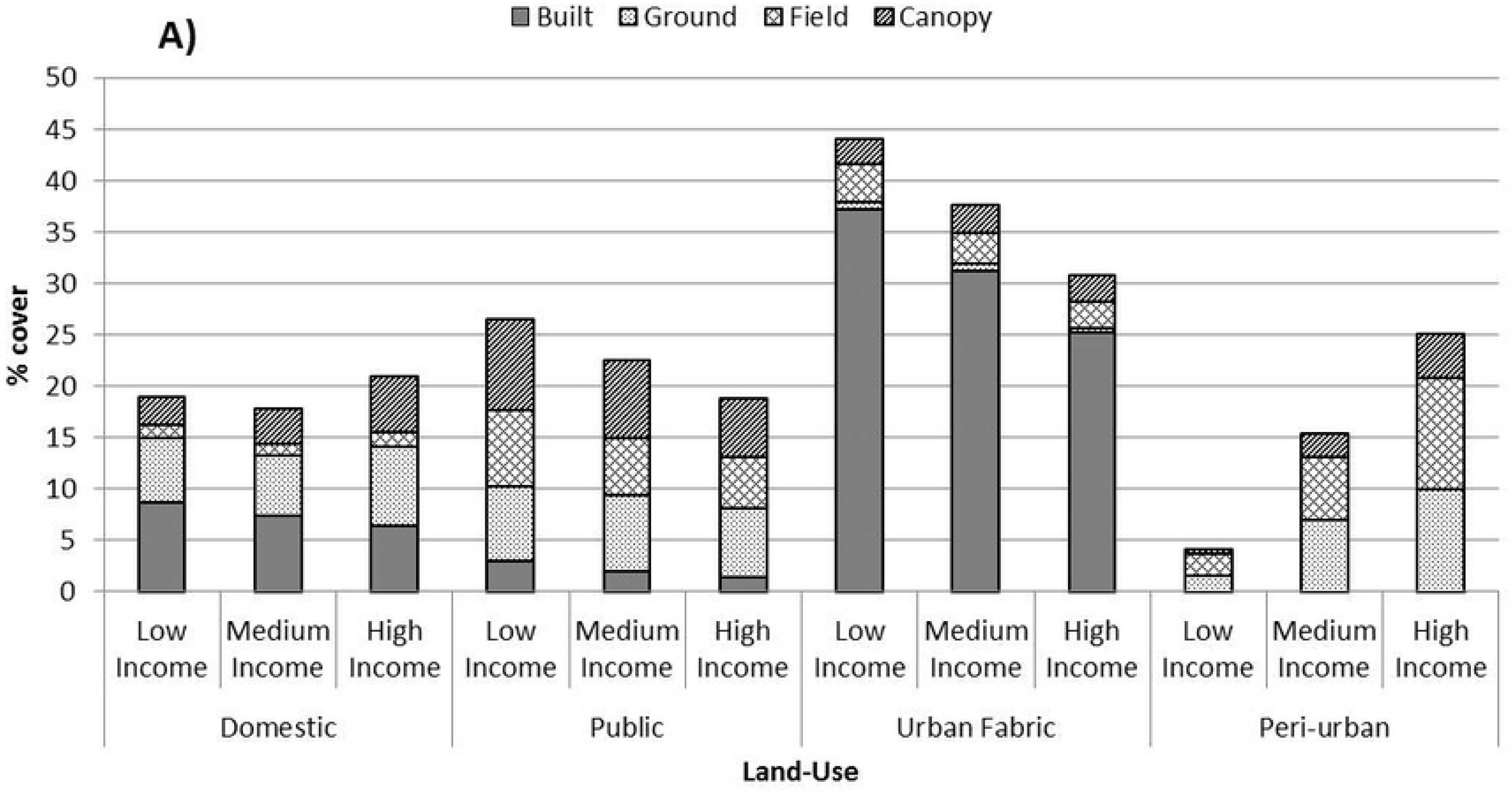
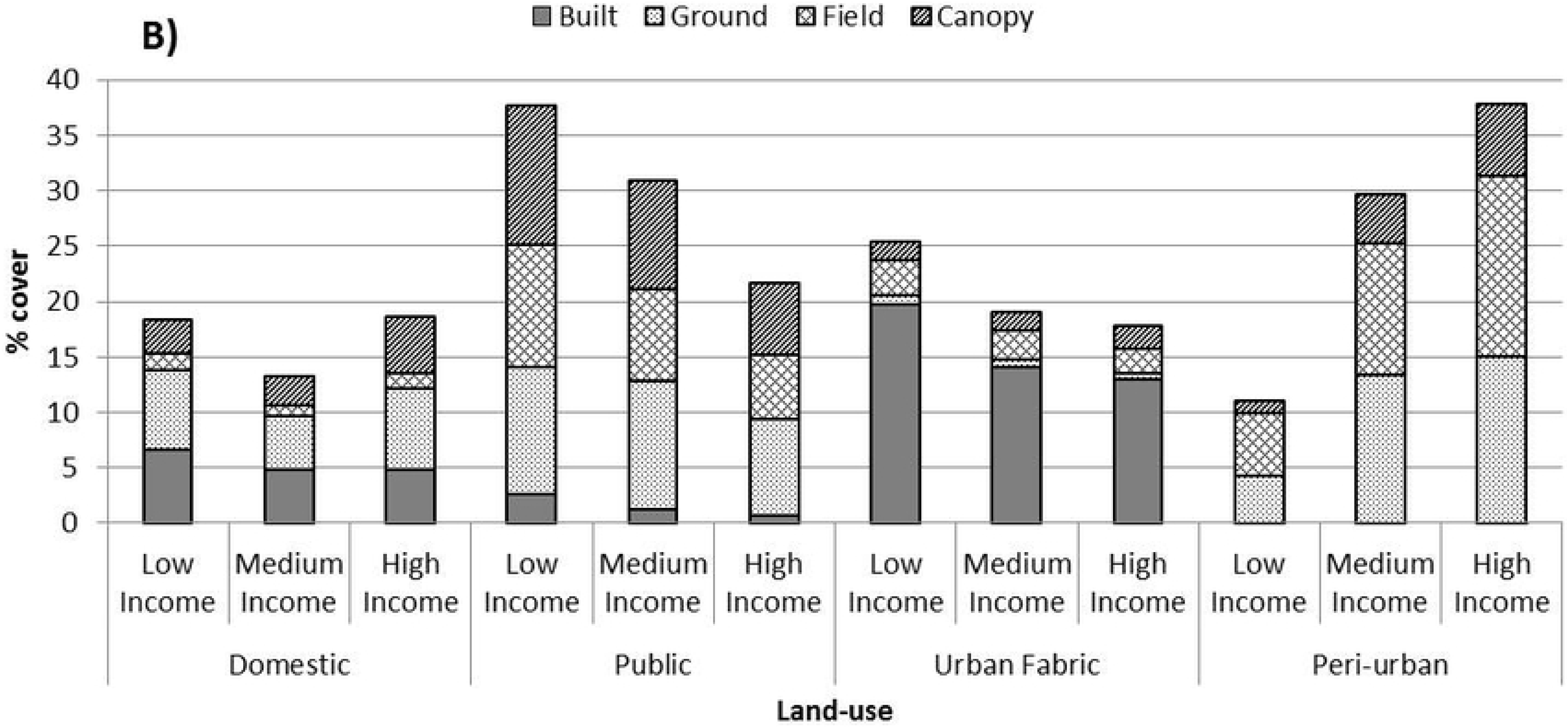
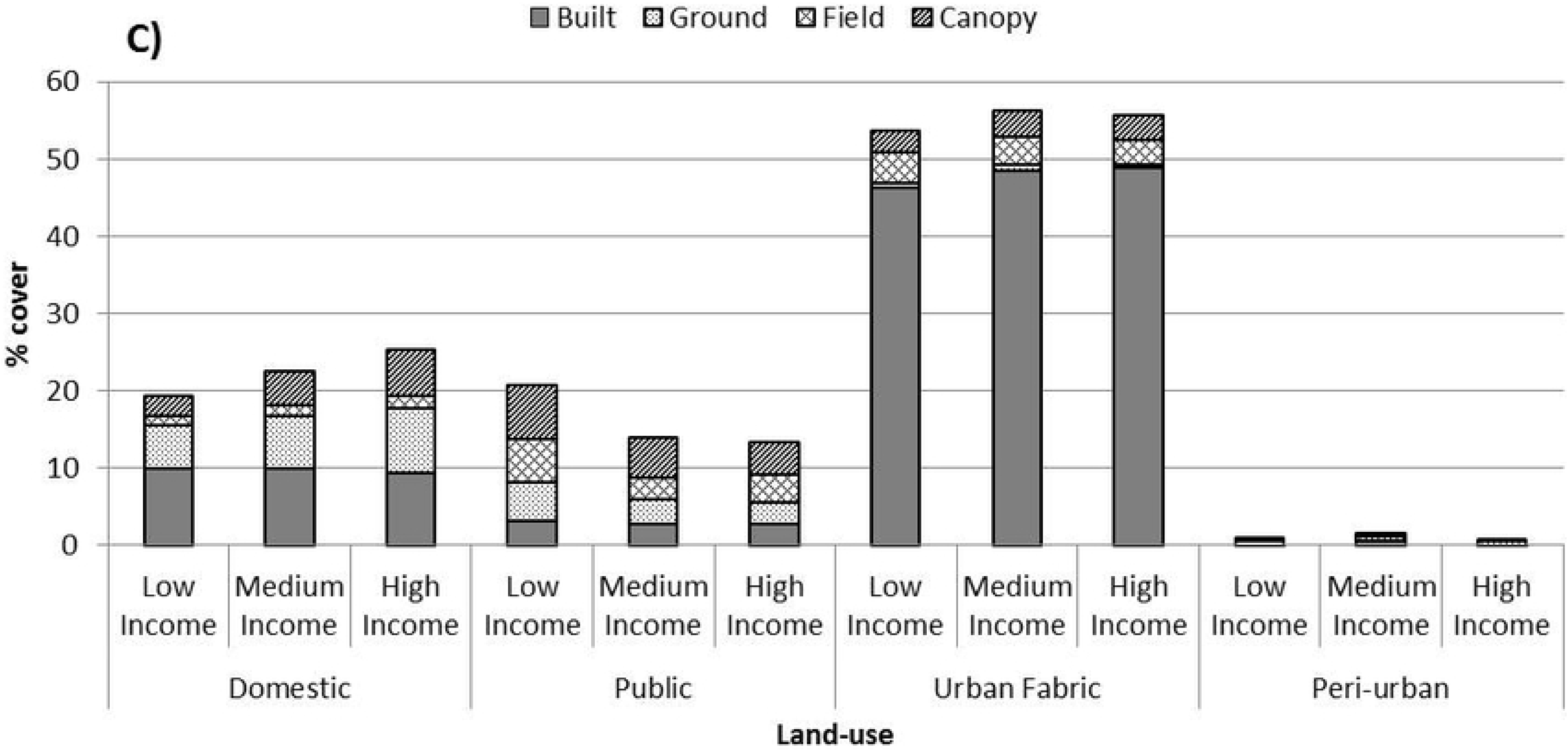
Vegetation cover within major land-uses (those comprising > 1 % of the study area) all areas; B) low-urban areas; C) high-urban areas.

The spatial extent and content of public and domestic green space exhibited contrasting mean values between low and high urban areas. Values associated with domestic gardens in particular also showed considerable variation as a function of income. For example, in terms of domestic green-space, low-urban areas contained lower cover relative to high-urban areas and, within the context of the latter, higher income was associated with both a larger spatial extent and a greater proportion of green land-cover. For both levels of urbanity, lower income areas contained the greatest public green space cover with a higher degree of surface sealing seen for this land-use in the high-urban context. Table 2 gives correlation coefficients (Pearson’s r) between land-use types and key indicators of urbanisation.

**Table 2.** Correlations between land-use and urban indicators (at 1 km^2^)

| Green-space type | Low-urban |  |  |  |  | High-urban |  |  |  |  |
| --- | --- | --- | --- | --- | --- | --- | --- | --- | --- | --- |
|  | Minor Rd Density | Major Rd Density | Population Density | Buildings Density | Mean Building Size | Minor Rd Density | Major Rd Density | Population Density | Buildings Density | Mean Building Size |
| Domestic | 0.886** | -0.042 | 0.802** | 0.932** | -0.228** | 0.552** | -0.376** | 0.546** | 0.955** | -0.694** |
| Public | 0.023 | 0.140 | 0.053 | 0.014 | 0.016 | -0.493** | -0.126 | -0.455** | -0.401** | -0.114 |
| Institutional | 0.504** | 0.217* | 0.590** | 0.504** | -0.055 | 0.247** | -0.026 | 0.260** | 0.152 | -0.192* |
| Urban Fabric | 0.740** | 0.359** | 0.727** | 0.713** | 0.082 | -0.168 | 0.435** | -0.214* | -0.619** | 0.738** |
| Peri-urban | -0.725** | -0.213* | -0.710** | -0.726** | 0.064 | -0.311** | 0.108 | -0.252** | -0.237** | 0.268** |

\* significant at the $p < 0.05$ level
\*\* significant at the $p < 0.01$ level

The relative cover by major land-use types for three quantile groups of the Largest Patch Index metric within 1 km^2^ zones (low LPI = land-sharing; high LPI = land-sparing), controlling for overall green land-cover, is given in Figure 6.

**Figure 6.**
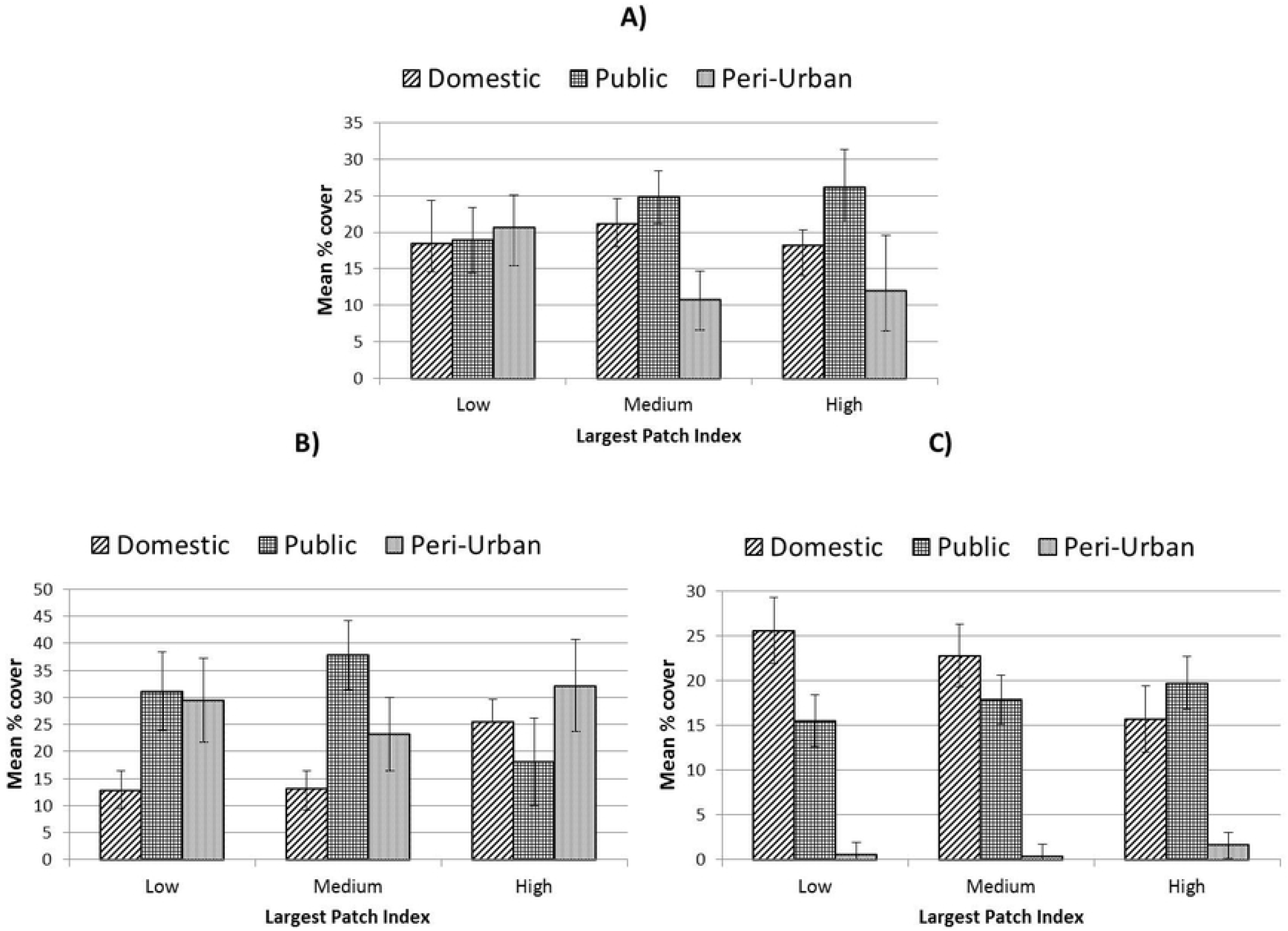
Relative extent of public, domestic and peri-urban green space at units of 1 km^2^ across a gradient of land sparing-sharing for A) all areas; B) low-urban areas and C) high urban areas. Error bars represent 95% confidence intervals.

Ecological and socio-environmental characteristics varied significantly as a function of land-sparing-sharing and urbanity. Figures 7, 8 and 9 and 10 give marginal mean values for TCA, Meff, SHDI and vNDVI respectively for low, medium and high quantile groups for LPI at 0.5, 1 and 2 km^2^.

**Figure 7.**
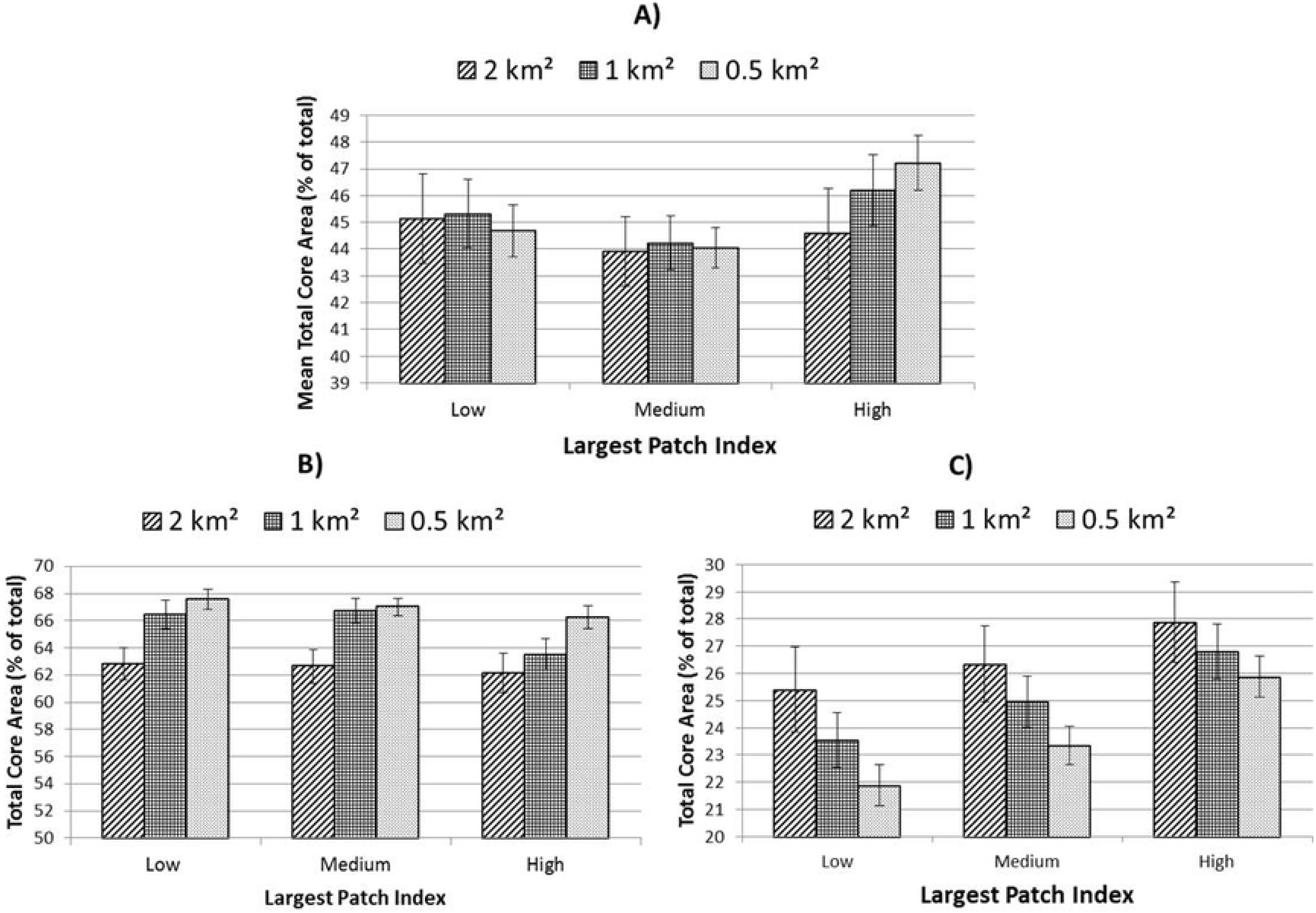
Mean Total Core Area for three levels of land-sparing/sharing controlling for overall green cover. A) all areas; low-urban areas and C) high urban areas. Error bars represent 95% confidence intervals.

**Figure 8.**
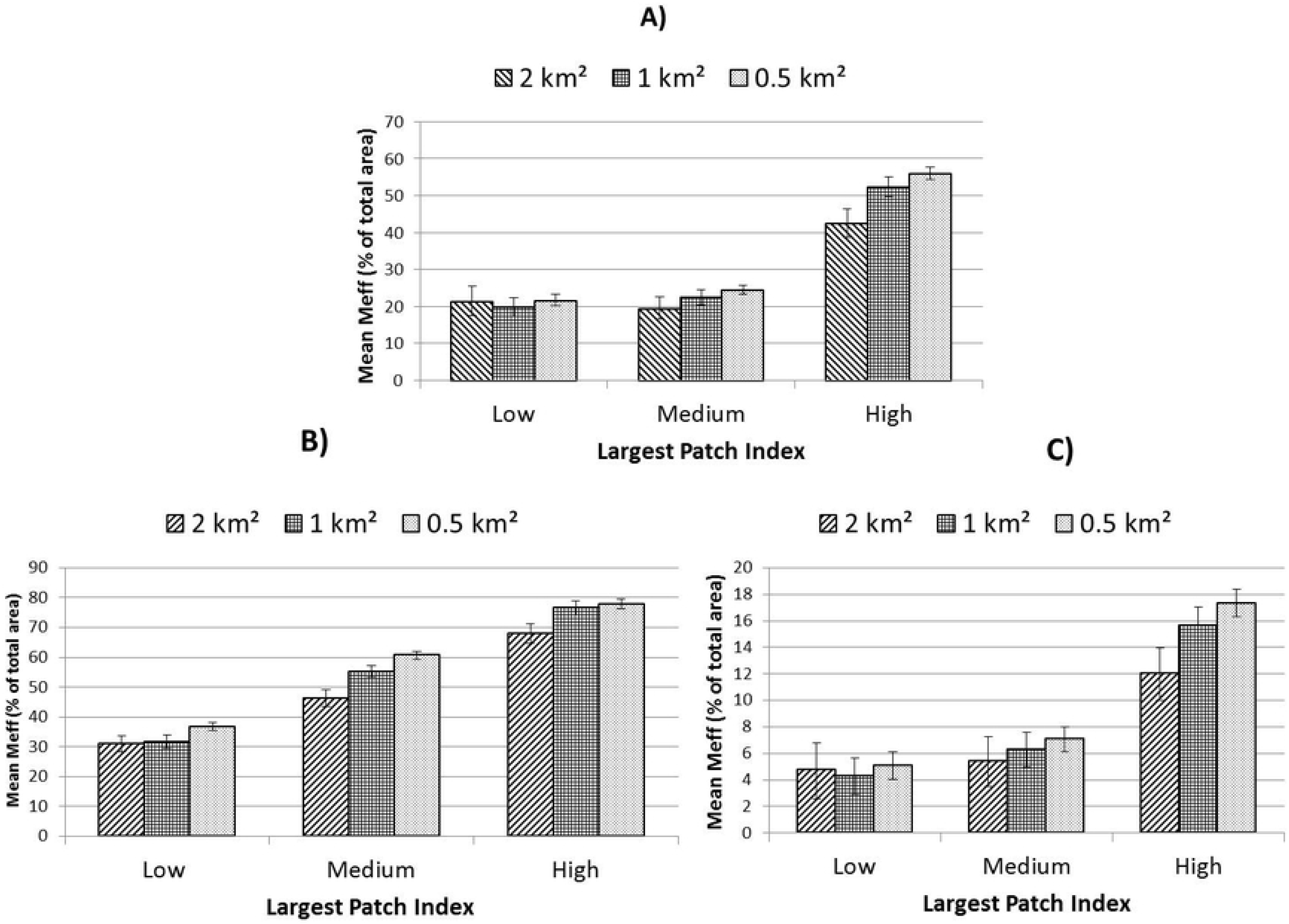
Effective mesh size for three levels of land-sparing/sharing controlling for overall green cover. A) all areas; B) low-urban areas and C) high urban areas. Error bars represent 95% confidence intervals.

**Figure 9.**
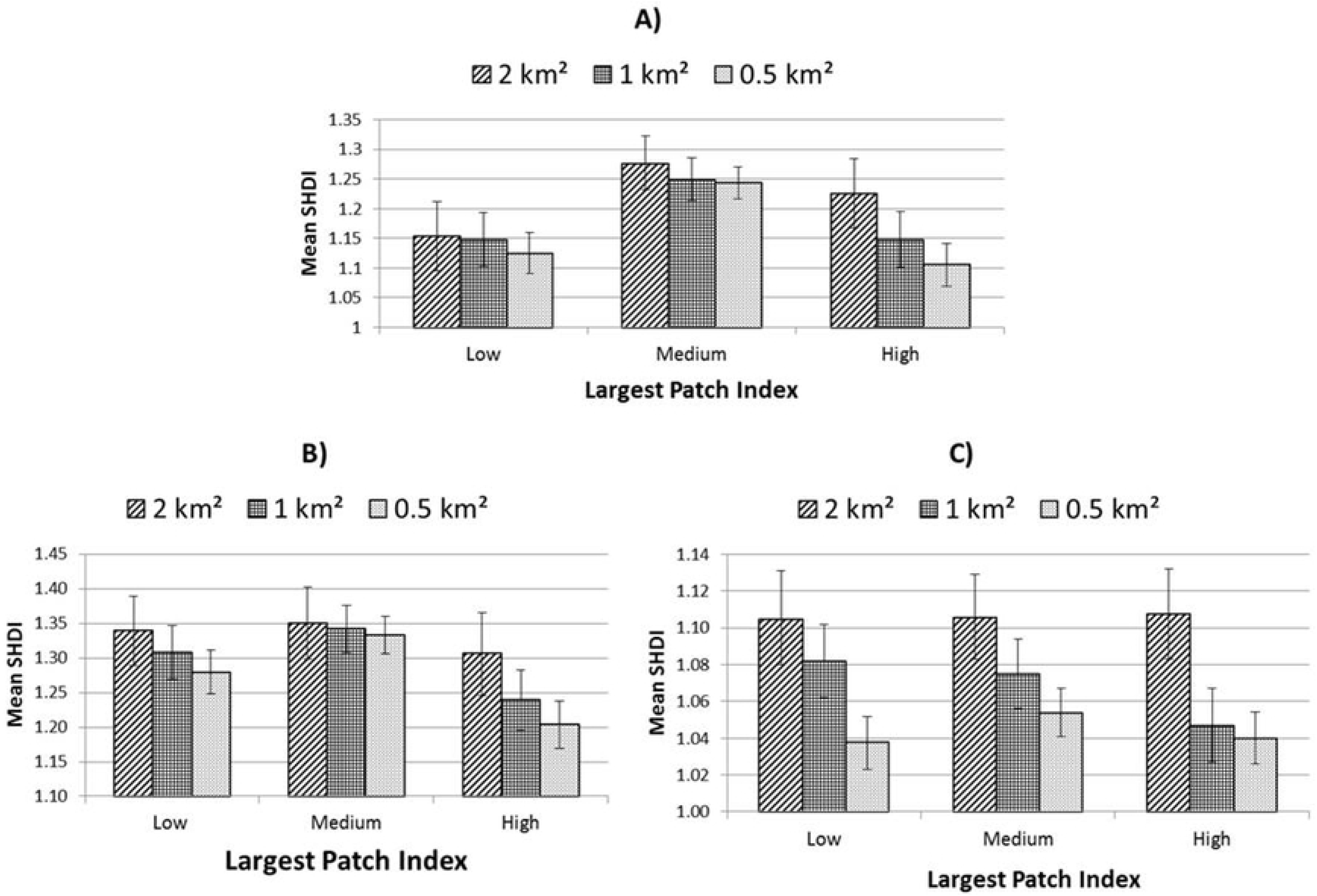
Mean SHDI for three levels of land-sparing/sharing controlling for overall green cover. A) all areas; B) low-urban areas and C) high urban areas. Error bars represent 95% confidence intervals.

**Figure 10.**
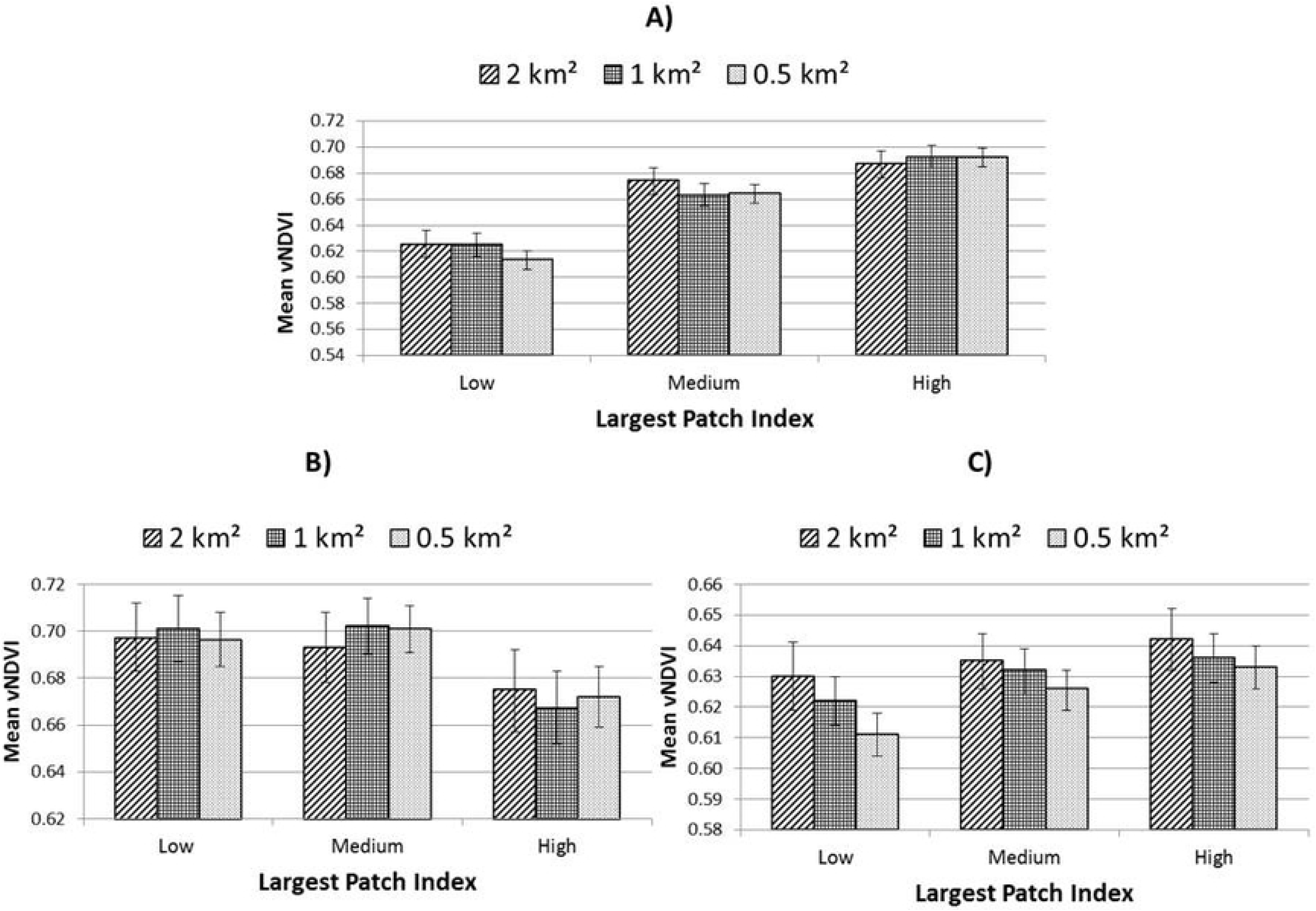
Mean vNDVI across three levels of land-sparing/sharing controlling for overall green cover. A) all areas; B) low-urban areas and C) high urban areas. Error bars represent 95% confidence intervals.

Contrasting patterns were observed between individual landscape metrics with TCA and SHDI in particular exhibiting unique distributions along the sharing-sparing gradient employed. Figures 11 and 12 give the marginal mean values resulting from general linear models for socio-environmental variables land surface temperature and ambient nitrogen dioxide concentration respectively. In terms of population within 300 m of a recreational green space, statistical significance was exhibited only in high urban areas (Figure 13)

**Figure 11.**
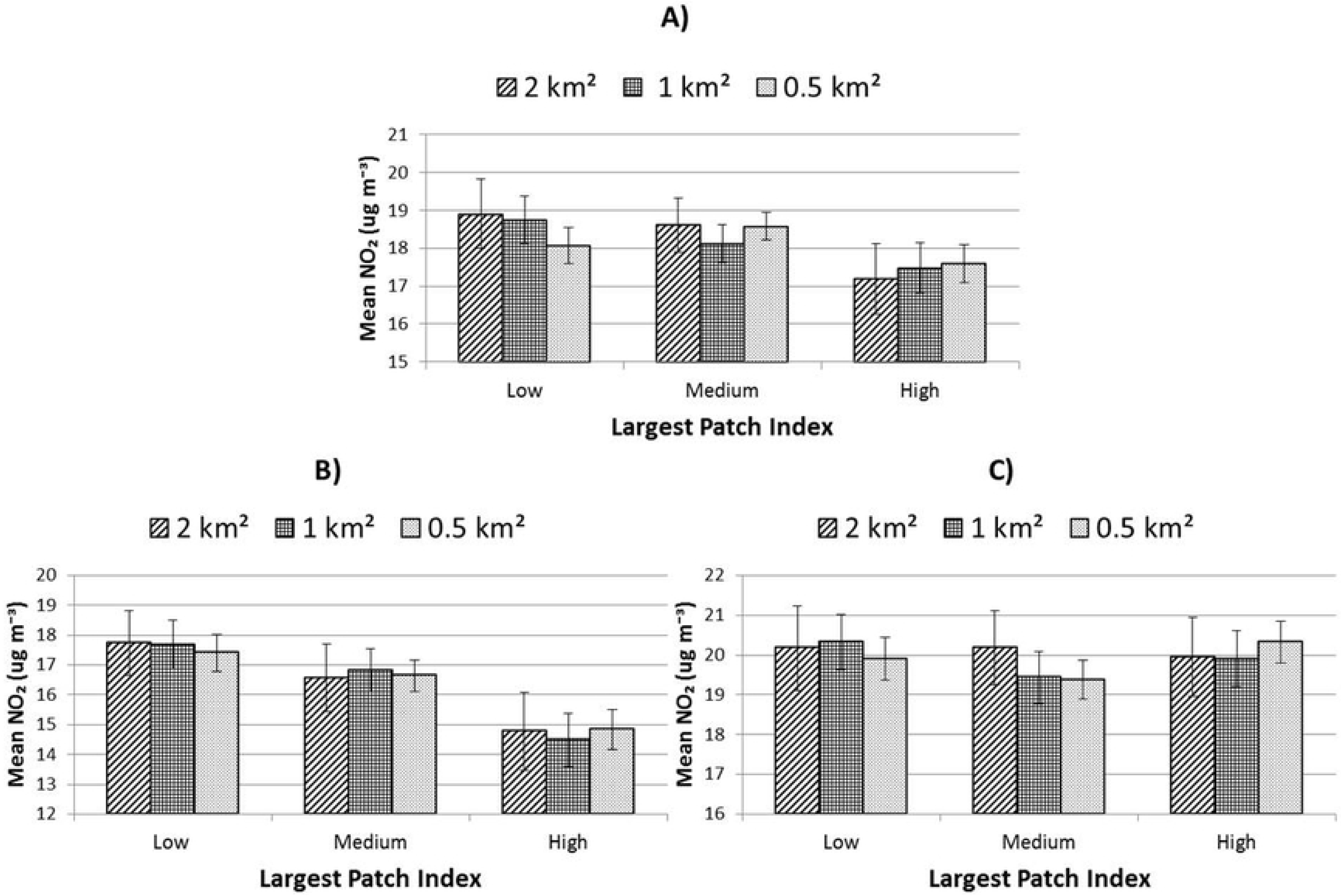
Mean ambient NO_2_-concentration for three levels of land-sparing/sharing controlling for overall green cover. A) all areas; B) low-urban areas and C) high urban areas. Error bars represent 95% confidence intervals.

**Figure 12.**
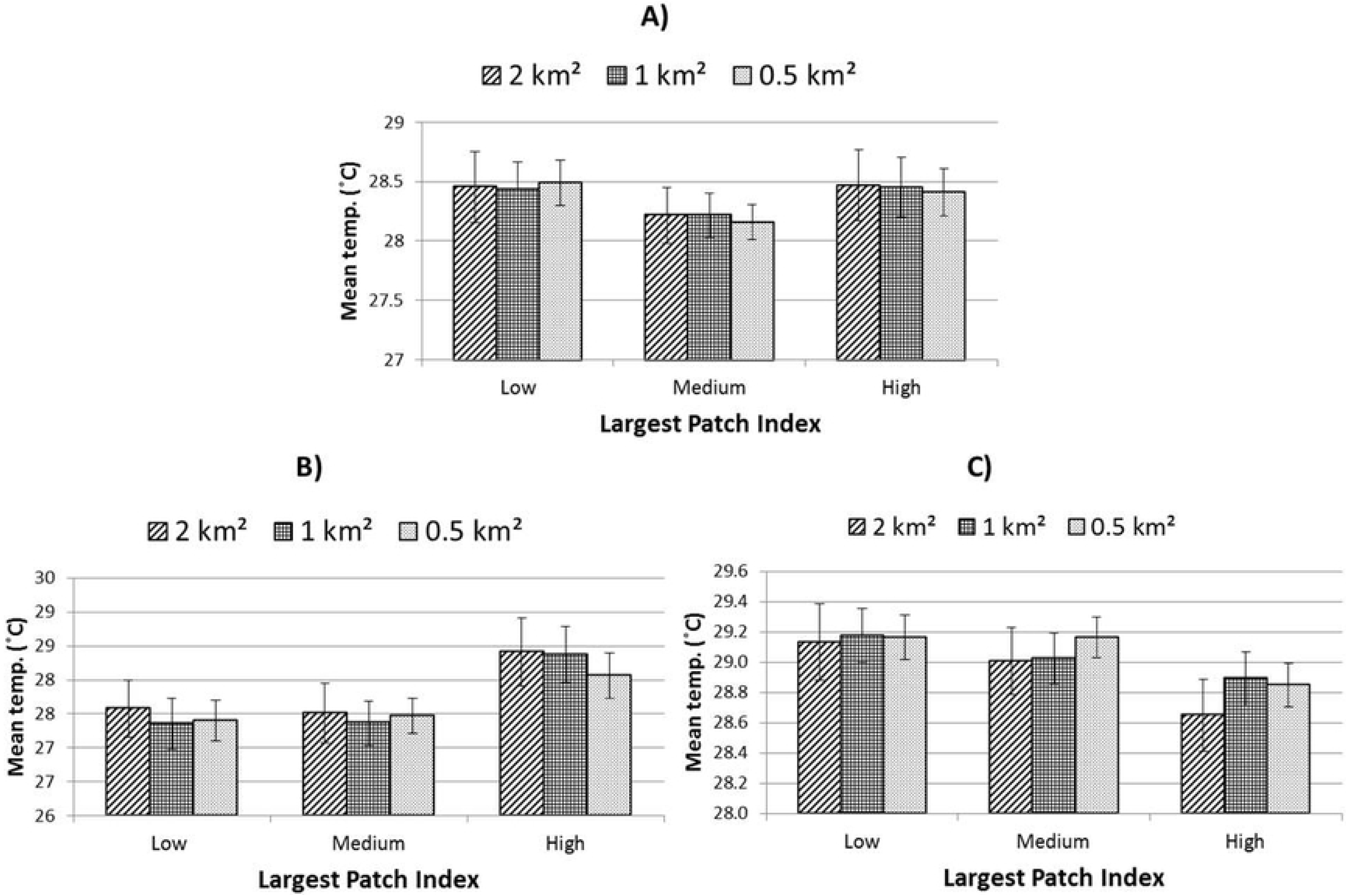
Mean land surface temperature for three levels of land-sparing/sharing controlling for overall green cover. A) all areas; B) low-urban areas and C) high urban areas. Error bars represent 95% confidence intervals.

**Figure 13.**
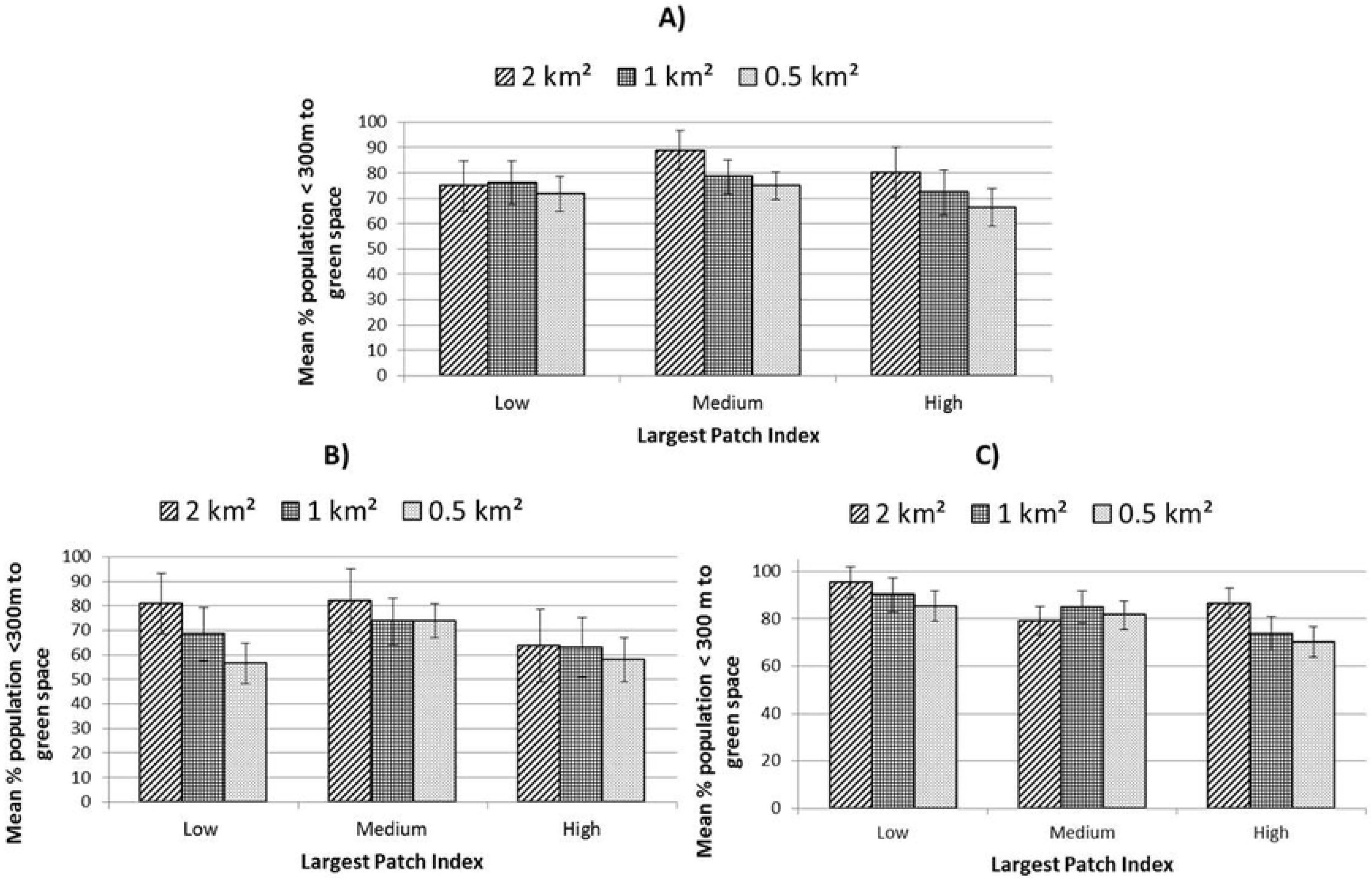
Mean percentage population within 300 m of a recreational green space across three levels of land-sparing/sharing controlling for overall green cover. A) all areas; B) low-urban areas and C) high urban areas. Error bars represent 95% confidence intervals.

Table 3 gives significance levels for models at each scale and level of urbanity considered. Overall, analyses at units of 0.5 km^2^ provided the greatest number statistically significant tests, though low-urban areas did not follow this trend as closely as high-urban areas.

**Table 3.** Significance levels (*p* values) for all general linear model analyses carried out in this study.

| Dependent variable | All areas |  |  | Low Urban |  |  | High urban |  |  |
| --- | --- | --- | --- | --- | --- | --- | --- | --- | --- |
|  | 0.5 km<sup>2</sup> | 1 km<sup>2</sup> | 2 km<sup>2</sup> | 0.5 km<sup>2</sup> | 1 km<sup>2</sup> | 2 km<sup>2</sup> | 0.5 km<sup>2</sup> | 1 km<sup>2</sup> | 2 km<sup>2</sup> |
| TCA | < 0.001 | 0.049 | 0.459 | 0.144 | <0.001 | 0.801 | < 0.001 | <0.001 | 0.100 |
| Meff | < 0.001 | < 0.001 | < 0.001 | < 0.001 | < 0.001 | <0.001 | < 0.001 | < 0.001 | < 0.001 |
| SHDI | < 0.001 | < 0.001 | 0.003 | < 0.001 | 0.003 | 0.617 | 0.163 | 0.050 | 0.991 |
| Mean Temp. | 0.005 | 0.160 | 0.234 | 0.020 | 0.002 | 0.040 | 0.003 | 0.108 | 0.025 |
| vNDVI | < 0.001 | <0.001 | 0.002 | 0.006 | 0.002 | 0.228 | < 0.001 | 0.072 | 0.301 |
| nitrogen dioxide | 0.004 | 0.070 | 0.045 | < 0.001 | < 0.001 | 0.007 | 0.033 | 0.187 | 0.936 |
| Pop. <300 m to green space | 0.164 | 0.558 | 0.054 | 0.001 | 0.391 | 0.203 | 0.007 | 0.009 | 0.004 |

### Multiple linear regression results

Table 4 gives the results of the multiple linear regression models with landscape metrics LPI, TCA, Meff, SHDI and vNDVI as dependent variables and Table 4 summarizes regression results where socio-environmental variables mean LST, mean nitrogen dioxide concentration and percentage population within 300 m of a recreational green space.

**Table 4.** Results of regressing land-use-land-cover attributes on landscape metrics used in this study. All tests carried out at 1 km^2^ units.

| Low-urban | Beta | Sig. | High-urban | Beta | Sig. |
| --- | --- | --- | --- | --- | --- |
| **LPI 1 km<sup>2</sup>** |  |  |  |  |  |
| r<sup>2</sup>: 0.64 |  |  | r<sup>2</sup>: 0.47 |  |  |
| Major road density | -0.510 | < 0.01 | Major road density | -0.228 | 0.002 |
| Domestic green cover | 0.321 | < 0.01 | Domestic green cover | 0.707 | <0.01 |
| Domestic built cover | -0.808 | < 0.01 | Domestic built cover | -0.689 | < 0.01 |
| Public built cover | -0.114 | 0.036 | Public green cover | 0.360 | < 0.01 |
|  |  |  | Peri-urban green cover | 0.180 | 0.008 |
| **TCA 1 km<sup>2</sup>** |  |  |  |  |  |
| r<sup>2</sup>: 0.89 |  |  | r<sup>2</sup>: 0.98 |  |  |
| Major road density | -0.169 | < 0.01 | Domestic built cover | -0.080 | < 0.01 |
| Domestic built cover | -0.874 | < 0.01 | Public green cover | 0.808 | < 0.01 |
| Public built cover | -0.284 | < 0.01 | Peri-urban green cover | 0.451 | < 0.01 |
| Peri-urban mean patch area | 0.96 | 0.002 | Public mean patch area | 0.058 | < 0.01 |
| Public green cover | 0.060 | 0.041 | Institutional green cover | 0.177 | < 0.01 |
|  |  |  | Domestic green cover | 0.596 | < 0.01 |
|  |  |  | Informal urban greenery | 0.210 | < 0.01 |
| **Meff 1 km<sup>2</sup>** |  |  |  |  |  |
| r<sup>2</sup>: 0.82 |  |  | r<sup>2</sup>: 0.67 |  |  |
| Domestic built cover | -0.808 | < 0.01 | Domestic built cover | -0.664 | < 0.01 |
| Major rd density | -0.458 | < 0.01 | Public green cover | 0.514 | < 0.01 |
| Domestic MPA | 0.160 | < 0.01 | Peri-urban green | 0.282 | < 0.01 |
| Public built cover | -0.224 | < 0.01 | Domestic green cover | 0.942 | < 0.01 |
| **SHDI 1 km<sup>2</sup>** |  |  |  |  |  |
| r<sup>2</sup> = 0.55 |  |  | r<sup>2</sup> = 0.92 |  |  |
| Peri-urban | -0.756 | < 0.01 | Informal Urban Greenery | 0.257 | < 0.01 |
| Informal Urban Greenery | 0.237 | 0.01 | Public green cover | 0.793 | < 0.01 |
| Domestic | -0.290 | < 0.01 | Domestic green cover | 0.712 | <0.01 |
|  |  |  | Public mean patch area | -0.067 | 0.029 |
|  |  |  | Peri-urban | 0.334 | < 0.01 |
|  |  |  | Institutional green cover | 0.210 | <0.01 |
| **vNDVI 1km<sup>2</sup>** |  |  |  |  |  |
| r<sup>2</sup>: 0.64 |  |  | r<sup>2</sup>: 0.75 |  |  |
| Public | 0.393 | < 0.01 | Domestic field | 0.251 | < 0.01 |
| Domestic built cover | -0.281 | < 0.01 | Domestic canopy | 0.360 | < 0.01 |
| Public built | -0.134 | 0.024 | Public field | 0.252 | < 0.01 |
| Public canopy | 0.241 | < 0.01 | Public canopy | 0.399 | < 0.01 |
| Institutional field layer | - | - | Institutional field layer | 0.112 | 0.018 |
| Peri-urban canopy | 0.513 | < 0.01 | Public built cover | -0.137 | 0.013 |
| Domestic mean patch area | 0.167 | < 0.01 | Major road density | -0.112 | 0.027 |
| Public mean patch area | 0.141 | 0.013 | Public mean patch area | 0.166 | < 0.01 |
| Peri-urban mean patch area | -0.367 | < 0.01 | Public ground | 0.226 | < 0.01 |

Regression analyses demonstrated that public and private land-uses exhibited unique and contrasting associations with ecological and socio-environmental variables implying considerable potential trade-offs. Moreover, these associations varied as a function of the level of urbanity and appeared to be modified by patch characteristics (mean area and green land-cover).

## Discussion

### Land-use characteristics and sharing-sparing scenarios

For the study area as a whole, and in areas of high urbanity, the distribution of public versus private green-spaces, controlling for total green land-cover, exhibited patterns that fulfill expectations of land-sparing-sharing scenarios. Inverse trends were observed for mean cover of public relative to domestic green space with increasing LPI (Figure 6a and c). However, in areas of low urbanity this pattern was not replicated where a dominance of public over domestic land-use was seen in land-sharing areas (i.e. low LPI) with domestic green space cover highest in land-sparing areas. Our analysis suggests, therefore, that the definition of land-sparing and sharing within an urban planning framework, in terms of primary land-uses which support this dichotomy, is subject to some fluidity as a function of urbanity. Moreover, the regression results highlighted domestic green and built land-covers as critical factors contributing to the largest patch index in both low and high urbanity areas, seemingly exerting a stronger influence on LPI than public green-space (Table 4). This is an important observation as it challenges some of the assumptions surrounding the relative patterns resulting from the prevalence of public and private green spaces within green infrastructure planning frameworks (Lin and Fuller, 2013). That ratios of built-to-green land-cover in domestic green space were also shaped by socio-economic status (Figure 5) suggests that overall urbanity, land-cover and economic status may all comprise determinants of land-sparing-sharing configurations in city regions.

### Level of Urbanity

Our analysis suggests that complex trade-offs may be implied by the ascendency of one or other of a land-sparing versus land-sharing approach within different contexts of urbanisation. This appeared to be most evident for socio-environmental factors considered. For example, models for mean LST and nitrogen dioxide values exhibited differing trends between high and low areas of urbanity. For mean LST, contrasting trends were observed along the sparing-sharing gradient between low and high-urban areas. This mirrored similarly inverse trends for domestic green space cover, presenting the latter as a potential causal factor. In the case of percentage of the local population in close proximity to a recreational green space, analysis of high-urban areas suggested provision was greatest in land-sharing environments when measured at a scale of 2 km^2^. For low-urban areas however, a mixture of land-sharing and land-sparing exhibited the greatest delivery of green space access. Vegetation quality (vNDVI) also exhibited highest mean values within this scenario in statistically significant models in low-urban areas (0.5 and 1 km^2^) whereas the highest values were associated with land-sparing in high-urban areas.

Although the two levels of urbanity presented some contrasting results, there was evidence of some consistency related to specific spatial or class-level components. For example, regardless of scale or level of urbanity, land-sparing appeared to consistently promote greater connectivity (Meff). That Meff was highest in land-sparing scenarios in both urbanity contexts (even though this implied different land-use patterns) suggests that individual land-use types are a minor consideration relative to spatial characteristics when aiming at connectivity. In terms of land-cover, tree canopy consistently promoted greater cooling (lower mean LST) and greater vegetation vigour, regardless of land-use or urbanity. This implies that, as identified by others (e.g. Collas et al, 2017), restoration through afforestation may significantly support and mediate broader landscape considerations in the promotion of urban ecosystem services and their resilience. From the perspective of landscape heterogeneity, differences in SHDI were significant between sparing-sharing scenarios in low-urban areas at the 0.5 and 1 km^2^ scale. At these scales, areas which comprised neither sharing nor sparing configurations exhibited greatest land-cover diversity, with land-sharing areas also showing significantly greater mean SHDI values than land-sparing areas (Figure 9). In addition, in low-urban areas peri-urban land-use appeared to play a detrimental role in landscape heterogeneity (Table 4). Overall, therefore, our results point towards an increase in vegetation diversity and quality in areas character rised by peri-urban land-use through the introduction of more typically urban green space types (Figures 5, 6 and 9). In the high-urban context, all major green land-uses appeared to contribute to landscape heterogeneity (Table 4) suggesting that increases in green land-cover of any type are beneficial regardless of land-sparing-sharing considerations (which were not statistically relevant to SHDI in high urban areas, Table 3).

### Scale

Associations between ecological and socio-environmental patterns and land-sparing-sharing scenarios appeared to be moderated as a function of the scale of investigation employed. For example, for the study area as a whole, when measured at units of 2 km^2^, TCA appeared to be highest within spatial configurations which represent land-sparing scenarios (Figure 7). In contrast, land-sparing appeared to promote this critical landscape characteristic when measured at scales of ≤ 1 km^2^. The influence of scale differed between variables. For example, of the landscape attributes tested, SHDI exhibited generally higher values when measured at larger scales, whereas (standardised) Meff values were highest at smaller scales of investigation. In terms of levels of statistical relevance, our analysis exhibited scale-dependence (Table 3). This is important from both an urban planning and nature conservation perspective. When treating the study area landscape as a whole, higher levels of statistical significance were exhibited at smaller scales of investigation for most variables considered (Table 3), though urbanity appeared to mediate this trend. For example, in low-urban areas, analysis at scale of 1 km^2^ returned the greatest number of statistically significant tests, whereas in high-urban areas this was occurred at the 0.5 km^2^ scale. This implies that in more highly fragmented landscapes, higher spatial resolution is necessary to discern land-sparing-sharing associations with environmental characteristics.

This variance as a function of scale and urbanity poses a challenge for landscape analysis which would inform decisions on social and ecological goals respectively. For example, analyses of species distributions in urban ecological studies are commonly carried out at units of 1 x 1 km^2^ (Vanbergen et al., 2005; Ockinger et al., 2009) though our results suggest that working at such scales may not capture the potential for land-cover configurations to similarly achieve co-benefits such as urban cooling. Therefore, using a multi-scale approach such as that developed here, considering multiple socio-environmental characteristics relevant to sustainable urban development may be of considerable merit. This is largely due to the possibility, as demonstrated here, of identifying optimum scales of analysis through relatively rapid assessments using GIS and remote sensing techniques.

#### Influence of land-cover

Regression analyses of individual land-use and land-cover attributes on environmental and ecological variables demonstrated a high degree of consistency between areas of contrasting urbanity though exceptions, related to SHDI in particular, were observed (Table 4). Specifically, both peri-urban and domestic land-use exhibited contrasting directions of association with SHDI dependent on whether they were assessed at low or high-urbanity. The cover by, and level of vegetation within, domestic gardens in particular were also subject to stark contrasts between areas of low and high urbanity (Figure 5). These disparities appeared to be underpinned by socio-economic processes. The latter, therefore, proved also to be an important local consideration moderating the status, and therefore influence, of land-use-land-cover combinations on ecological and environmental variables.

Cover by gardens and land-cover within gardens exhibited strong links with all socio-environmental characteristics measured. Of all land-cover types, mean LST was most strongly (negatively) associated with canopy cover in gardens in high-urban areas (Table 5), suggesting that management of domestic greening presents opportunities for climate resilience in cities. Green land-cover within informal and other private (institutional) settings also exerted significant influence on both ecological and environmental characteristics, particularly in high urban areas. This underlines the complex mosaic of land-uses contributing to effective urban green infrastructure and the need for land management within such spaces to be acknowledged as key components of planning for sustainable and resilient cities. Gardens also appeared to exert an influence on both proximity to green space and air quality. For example, domestic garden cover was positively associated with access to green space in high-urban areas though, notably, public green-space (to which category green recreational spaces belonged), was non-significant. This suggests that, for the current study area at least, access (defined as proximity) to recreational green spaces may be more closely related to population distribution than to provision of green space *per se*. This is supported by the fact that domestic green space mean patch size – denoting lower housing (and therefore population) density -was negatively associated with proximity to recreational green space (Table 5). This pattern supports other work on urban land-sparing which highlights the merits of land-sharing configurations on green space use (Soga et al., 2015). It also suggests, however, that increasing urban residential density, through compaction and in-filling may offer opportunities for sparing non-developed land whilst ensuring local access to green space.

**Table 5.** Results of regressing land-use-land-cover attributes on socio-environmental metrics used in this study. All tests carried out at 1 km^2^ units.

| Low-urban | Beta | Sig. | High-urban | Beta | Sig. |
| --- | --- | --- | --- | --- | --- |
| Mean LST |  |  | Mean LST 1 km<sup>2</sup> |  |  |
| r<sup>2</sup> = 0.68 |  |  | r<sup>2</sup> = 0.67 |  |  |
| Public ground | 0.311 | < 0.01 | Urban water | -0.324 | < 0.01 |
| Urban water | -0.182 | < 0.01 | Major road density | -0.215 | < 0.01 |
| Minor road density | 0.375 | < 0.01 | Public canopy | -0.338 | < 0.01 |
| Public canopy | -0.425 | < 0.01 | Informal Urban Greenery mean patch area | -0.405 | - |
| Peri-urban canopy | -0.632 | < 0.01 | Public field layer vegetation | -0.264 | < 0.01 |
| Informal Urban Greenery | -0.162 | 0.19 | Domestic canopy | -0.529 | < 0.01 |
| Peri-urban mean patch area | -0.160 | 0.013 | Institutional canopy | -0.206 | 0.027 |
| Peri-urban mean patch area | 0.187 | < 0.01 | Domestic mean patch area | -0.295 | < 0.01 |
| Public mean patch area | -0.125 | 0.022 | Public water | -0.109 | < 0.01 |
| Domestic canopy | -0.210 | < 0.01 |  |  |  |
| Public field layer vegetation | -0.265 | < 0.01 |  |  |  |
| Nitrogen dioxide |  |  |  |  |  |
| r<sup>2</sup> = 0.59 |  |  | r<sup>2</sup> = 0.66 |  |  |
| Major road density | 0.259 | < 0.01 | Major road density | 0.382 | < 0.01 |
| Peri-urban field layer | -0.496 | < 0.01 | Peri-urban mean patch area | -0.184 | < 0.01 |
| Public canopy | 0.274 | < 0.01 | Institutional built | 0.234 | < 0.01 |
| Domestic mean patch area | -0.200 | < 0.01 | Domestic green cover | -0.465 | < 0.01 |
| Public field layer | -0.208 | < 0.01 | Institutional field layer | -0.234 | < 0.01 |
| Buildings density | 0.147 | 0.016 | Informal Urban Greenery | 0.223 | < 0.01 |
|  |  |  | Minor road density | 0.332 | < 0.01 |
| Pop < 300 m green space |  |  |  |  |  |
| r<sup>2</sup> = 0.63 |  |  | r<sup>2</sup> = 0.54 |  |  |
| Peri-urban green cover | -0.545 | < 0.01 | Domestic | 0.739 | < 0.01 |
| Major road density | 0.211 | < 0.01 | Minor road density | 0.453 | < 0.01 |
| Peri-urban mean patch area | -0.160 | 0.013 | Informal Urban Greenery | 0.307 | < 0.01 |
| Informal Urban Greenery mean patch area | -0.198 | 0.01 | Domestic green cover | -0.218 | < 0.01 |
| Public mean patch area | -0.146 | 0.09 | Institutional green cover | 0.157 | 0.018 |
|  |  |  | Informal Urban Greenery mean patch area | -0.391 | < 0.01 |

In terms of air quality, domestic garden cover showed a surprising negative association with mean nitrogen dioxide concentrations: the strongest of all land-uses types for high urban areas. Specific land-covers within gardens did not seem to be responsible for this association (Table 5), but that garden cover correlated negatively (p < 0.01) with density of major roads (Table 2) may offer a potential explanation and suggests urban form, rather than land-cover, as a critical factor. This idea is supported by results reported elsewhere which suggest that complex geometric patterns created by fragmented urban forms may reduce traffic-related congestion and pollution (Zhou et al., 2018). That tree cover in public green spaces in low-urban areas was positively associated with mean nitrogen dioxide concentrations may explain to some degree why public green-space cover overall was not statistically relevant to mean nitrogen dioxide concentrations. This stands in contrast to findings in other studies highlighting the ability of trees to remove nitrogen dioxide from the environment (Fantozzi et al., 2015). However, ours is the first study of its kind to consider a range of vegetation types across different land-uses simultaneously. The results of our regression models showed that tree canopy and lower vegetation types exhibited contrasting associations with level of nitrogen dioxide with field layer vegetation showing the greatest negative influence on ambient nitrogen dioxide at both levels of urbanity. Broader evidence on the relationship between the urban canopy and ambient nitrogen dioxide is, however, mixed (Yli-Pelkonen et al., 2018) and known to be subject to meteorological factors (Grundström et al., 2015). Specifically, ambient nitrogen dioxide has been shown to decrease with local air temperature (Ibid.). The latter is particularly relevant given that tree cover was negatively associated with LST in our results and implies a potential trade-off resulting from different socio-environmental outcomes related to the presence of green infrastructure (i.e. urban cooling and air quality). Overall, cover by water in urban areas suggested the greatest cooling effect by any land-cover, underlining the importance of waterways and wetlands in the regulation of the urban micro-climate (e.g. Gomez-Baggethun et al., 2013).

### Moving the land-sparing-sharing debate forward in urban areas

The analysis presented here demonstrates how a landscape approach, incorporating spatially coincident measures of land-use and land-cover, can be employed to unpick spatial and ecological complexities relevant to sustainable urban development. Our analysis suggests three pathways for future evaluation and research on landscapes subject to the process of urbanization. Firstly, scale (spatial units) should be considered in planning and research where multiple socio-environmental concerns are to be addressed. In the case of the former, we suggest that a modular approach working at smaller, local scales of analysis should be employed to capture variables that are highly spatially sensitive. Concurrently, research should focus on evaluating the potential for up-scaling analysis of small-scale phenomena (e.g. micro-climate regulation) to align with larger theoretically established units of investigation of others (e.g. species distribution). Secondly, spatial context in terms of levels of urbanity should be equally considered as a highly significant mediating factor in the determination of optimal land-use configurations. Not only do levels of urbanization modify the spatial characteristics of landscapes, but from the perspective of landscape resilience and ecosystem services provision, different contexts will dictate the nature of management goals related to spatial planning. For example, in urban areas where natural green cover is high fragmented but may also exhibit high heterogeneity, developing landscape configurations which increase connectivity per unit area may take priority over increasing diversity. Conversely, in peri-urban areas where green cover consists of larger and more connected patches, but highly homogenous (e.g. due to agricultural practices), land-use-land-cover combinations which promote landscape complexity rather than cohesion may be prioritised. Further, our results suggests that, even when different landscape configurations are promoted in urban and peri-urban areas, this may in reality involve parallel promotion of the same land-use type. However, we concede that the current study used a highly simplified dichotomous take on an urban-to-peri-urban gradient, controlling for overall green land-cover within each zone. In reality urban-rural gradients will consist of multiple degrees of urbanisation and human density. Furthermore, overall greenness of the environment and the merits of land-sparing versus sharing outcomes are likely to be subject to non-linear functional relationships (Stott et al., 2015). Therefore, our findings should be tested, ideally across landscapes which exhibit multiple combinations of green land-cover and population, in order to identify potential thresholds in the relative performance of land-sparing-sharing combinations.

Land-use-land-cover combinations exerted a significant influence on the social-ecological-environmental characteristics explored here and exhibited the potential to subvert assumptions related to land-sparing-sharing scenarios (e.g. the relative distribution of public and private green space). We suggest, therefore, as a third imperative for future research on land-use configurations towards sustainable urban landscapes, that land-cover specifically (and ecological restoration more broadly) be embedded within research designs as a qualitative consideration with a view to potentially clarifying and resolving tensions related to spatial considerations. Operationalising and refining these three principles of analysis could help to clarify and harness complexity in human-dominated landscapes towards spatial configurations that promote productive, diverse and ultimately resilient urban areas

